# Virgin Birth: A genetic basis for facultative parthenogenesis

**DOI:** 10.1101/2022.03.13.484157

**Authors:** A. L. Braun, D. K. Fabian, E. Garrison, D. M. Glover

## Abstract

Sexual reproduction evolved 1-2 billion years ago and underlies the biodiversity of our planet. Nevertheless, devolution of sexual into asexual reproduction can occur across all phyla of the animal kingdom. The genetic basis for how parthenogenesis can arise is completely unknown. To understand the mechanism and benefits of parthenogenesis, we have sequenced the genome of the facultative parthenogen, *Drosophila mercatorum*, and compared its organisation and expression pattern during parthenogenetic or sexual reproduction. We identified three genes, *desat2*, *Myc*, and *polo* in parthenogenetic *D. mercatorum* that when mis-regulated in a non-parthenogenetic species, *D. melanogaster*, enable facultative parthenogenetic reproduction. This simple genetic switch leads us to propose that sporadic facultative parthenogenesis could evolve as an ‘escape route’ preserving the genetic lineage in the face of sexual isolation.

## Introduction

Parthenogenesis is a form of reproduction resulting in uniparental offspring having only the maternal genome; it is a virgin birth. There are two types of parthenogenesis: facultative, having the ability to switch back to sexual reproduction; and obligate, in which this is not possible. Sexual reproduction requires a carefully orchestrated program whereby the genome is first duplicated before undergoing two divisions in the absence of DNA synthesis to generate a complement of haploid gametes that can be combined with those of the opposite sex, or mating type in the context of lower eukaryotes, to generate a diploid zygote. Facultative parthenogens retain the key meiotic machinery and yet have a hitherto unknown, but likely heritable, change that enables them to regain diploidy after meiosis and initiate mitotic divisions. By contrast, obligate parthenogens can theoretically have a block anywhere in meiosis and may eliminate it completely. It is therefore likely that different mechanisms underlie parthenogenesis depending upon which stage of sexual reproduction is blocked.

Parthenogenesis was first observed in aphids by Charles Bonnet in approximately 1740 and yet, its underlying mechanism has not been identified in any animal. Despite being poorly understood, parthenogenesis is generally regarded as being a deleterious reproductive strategy because it fails to generate genetic diversity. Nevertheless, parthenogenesis has evolved repeatedly across different phyla of animals and plants. One reason for the failure to identify any genetic cause of naturally occurring parthenogenesis in animals is that ancient obligate parthenogenetic lineages are often compared to similar, sexually reproducing counterparts that have sometimes diverged millions of years ago. It then becomes impossible to separate the primary cause from multiple downstream consequences. If we are to understand parthenogenesis, we must look at new species or, better yet, examine those able to switch from sexual to parthenogenetic reproduction. We postulated that a genetic cause likely underlies facultative parthenogenesis because it can undergo selection in *Drosophila*, locuts, and chickens and increase in frequency over several generations [1–4]. We therefore sought to uncover the genetic cause behind facultative parthenogenesis in *Drosophila mercatorum*,by sequencing its genome and comparing gene expression patterns during the oogenesis of females undertaking sexual or parthenogenetic reproduction. We now report a genetic cause of sporadic facultative parthenogenesis uncovered in *D. mercatorum* and show how these traits can be transferred to a sexually reproducing species, *Drosophila melanogaster*.

## Results

### The parthenogenetic ability of D. mercatorum

The facultative parthenogen, *D. mercatorum*, is unique in that some strains can behave as obligate parthenogens upon transitioning to parthenogenetic reproduction and certain strains can then be maintained in the lab indefinitely as healthy and easily expandable female only stocks [4–6]. *D. mercatorum* belongs to the *repleta* species group of South American cactus feeders, which are approximately 47 My diverged from *D. melanogaster* [7]. However, *D. mercatorum* appears invasive and has spread, far beyond the range of most other *repleta*, to Australia and as far north as New York [4, 8]. As nearly all strains of *D. mercatorum* studied to date show some degree of parthenogenetic capability [4], we began by determining the baseline of parthenogenesis in 8 different *D. mercatorum* strains using a classical assay adapted from the first study of *Drosophila* parthenogenesis [3]. Large numbers of virgin females were maintained on fresh food for the duration of their lives and the food examined for offspring at any developmental stage. The numbers of progeny ranged from the generation of a small number of developing embryos that died before hatching to the production of a small number of fertile adult flies (Data table S1). We observed that parthenogenetic offspring were produced on average halfway through the maximum life of the mother (Data table S1), at 22.2 days of the 50-day maximum lifespan. We also confirmed by PCR that these lines did not carry any Wolbachia infection (Data table S1), which is known to cause parthenogenesis in other arthropods [9], although Wolbachia is only reported to cause cytoplasmic incompatibility in *Drosophila* [10]. We also confirmed that the strains examined were indeed all *D. mercatorum* since they were able to interbreed producing viable and fertile male and female offspring, although the parthenogenetically reproducing strain had slight impediment to breeding and did not consistently produce offspring (Data table S2). As a result of these experiments, we selected two *D. mercatorum* stains for further study, a parthenogenetic strain from Hawaii and a sexually reproducing strain with very low parthenogenetic capability from São Paulo, Brazil.

### The genome of D. mercatorum

In search for genetic changes permitting parthenogenesis, we chose to sequence and compare the genomes of the chosen sexually reproducing and parthenogenetic strains of *D. mercatorum*. We produced polished chromosome-level genome assemblies, using Oxford Nanopore Technology (ONT) and Illumina sequencing technology. The quality of the assemblies was assessed using standard metrics of N50, coverage, and genome size (Table 1), all of which indicated that the genomes were of similar or greater quality to other *de novo Drosophila* genomes found on public databases. The genomes were then annotated using the *D. melanogaster* homologues, and we found that the sexually reproducing genome was slightly more functionally complete (Table 1). Most of the genes were on the 14 largest contigs of the parthenogenetic genome assembly but were distributed to a more non-linear degree across more contigs on the sexual genome assembly (Fig. S1A-B). This is likely a consequence of the positioning of more repetitive sequences in the sexual genome assembly (Table 1). We ensured that the sequencing depth and coverage were uniform by plotting the reads over the assembled genome (Fig. S1C-D). Together, these analyses indicated that both *de novo D. mercatorum* genome assemblies were of similar high quality.

**Table 1:** Analysis of the sexual and parthenogenetic *D. mercatorum* genomes. Genome assembly data metrics, quality control, and annotation metrics.

| Metric |  | Sexual | Parthenogenetic |
| --- | --- | --- | --- |
| Genome size |  | 171,182,504bp | 161,570,079bp |
| Coverage |  | 88x | 96x |
| Contig number |  | 556 | 330 |
| Continuity | Mean contig Length | 307,882bp | 489,606bp |
|  | Scaffolds NG50 | 22,671,956bp | 16,356,382bp |
| Repeat Content | Bases in repetitive regions | 34,751,631bp | 29,808,408bp |
|  | Percent of genome repetitive | 20.30% | 18.45% |
|  | GC Content | 40.65% | 40.33% |
| Functional completeness | Genes | 19,983 | 17,611 |
|  | Uniquely mapped reads (STAR) | 80-91% | 74-89% |

Since the first *Drosophila* genome sequenced was *D. melanogaster*, and due to its extensive adoption as a model organism, it is the benchmark *Drosophila* reference genome. *D. melanogaster* has four pairs of chromosomes, which in relation to their centromeres have 6 chromosome arms. These arms, referred to as the Muller elements A-F [11], correspond to the respective X-chromosome, the left and right arms of chromosomes 2 and 3, and chromosome 4. We found 24.4% divergence between both sexually (Fig. 1A) and parthenogenetically (Fig. S2A), reproducing *D. mercatorum* genome assemblies and the *D. melanogaster* reference genome (release 6). There was clustering of each contig from the two *D. mercatorum* genomes to specific chromosome arms in *D. melanogaster*, indicative of the shuffling of genes, which largely remain on the same chromosome arms. We also confirmed the matching of contigs to chromosome arms by checking the DNA k-mers using Nucmer (Fig. S3A-B). Our findings accord with long-held knowledge of how corresponding Muller elements form a series of homologous genetic ‘building blocks’ in different *Drosophila* species within which synteny is lost [11–13].

**Fig. 1.**
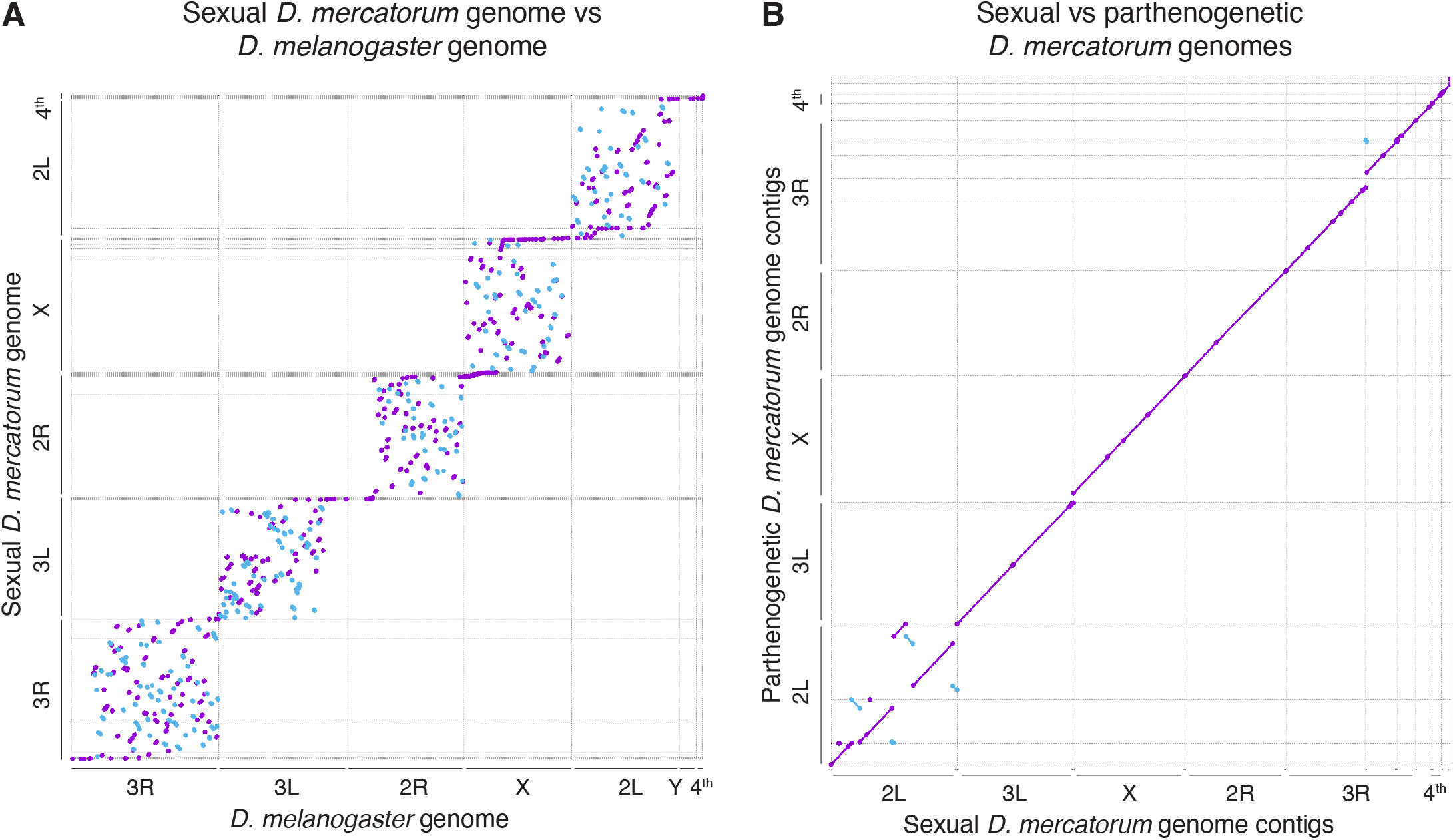
Sexual and Parthenogenetic *D. mercatorum* genome comparisons. (**A**) Sexual *D. mercatorum* genome assembly aligned against the *D. melanogaster* reference genome (release 6). In both genome comparisons the purple dots or lines represent sequences matching against the forward strand and the blue against the reverse. (B) Alignment of the parthenogenetic *D. mercatorum* genome against the sexual genome.

Together these analyses indicated that the chromosome-level genome assemblies for the sexually and parthenogenetically reproducing *D. mercatorum* strains were suited to detailed comparison between each other and with the *D. melanogaster* genome. When aligned, the sexual and parthenogenetic genomes were highly similar having only 1.2% divergence (Fig. 1B), which is consistent with pairwise heterozygosity, and thus further confirming that they are indeed the same species. Our comparison also revealed inversions on the 2L chromosome arm (Muller element B) which had previously been noticed between *D. mercatorum* populations collected from South and North America [8, 14]. These inversions are similar to that found in different populations of *Drosophila pseudoobscura* [15]. Therefore, these inversions may be instrumental in maintaining isolation between the different populations or simply a gauge of isolation and the initial stages of speciation.

We next confirmed that the genome assemblies matched the karyotypes of the sexual and parthenogenetic of *D. mercatorum* strains by localising local sequence markers onto preparations of mitotic chromosomes from *D. mercatorum* third instar larval brains using a fluorescence in situ hybridization (FISH) hybridisation chain reaction (HCR) protocol that we developed for this purpose (Supplementary text). We selected single genes within syntenic blocks that are conserved between *D. melanogaster* and *D. mercatorum* to serve as markers for each of the 6 chromosome arms of the mitotic karyotype (Fig. 2A). This allowed us to identify the Muller elements, A-F, for both sexual and parthenogenetic *D. mercatorum* strains (Fig. 2B). We found the fusion of the 2L/B and 3R/E (*D. melanogaster/Muller)* chromosome arms that was previously documented as unique to *D. mercatorum* within the *repleta* group [14] and the remaining chromosome arms were telocentric. We also observed that the 4^th^ chromosome of the parthenogenetic strain was substantially larger than the 4^th^ chromosome of the sexually reproducing strain. We also physically positioned the 14 largest contigs from the parthenogenetic genome onto the 3^rd^ instar larval salivary gland polytene chromosomes of both the sexually reproducing and parthenogenetic strains of *D. mercatorum* also using HCR FISH (Fig. 2C-D). Each contig mapped to the chromosome arm as predicted by the annotation and nucleotide sequence but the 4^th^ (F) chromosome of sexually and parthenogenetically reproducing strains of *D. mercatorum* were of similar size. This suggests that the increased size of the 4^th^ chromosome in diploid cells of the parthenogenetic strain is due to acquisition of satellite, heterochromatic sequences that do not undergo endoreduplication in the generation of polytene chromosomes. We conclude that the two chromosome-level genome assemblies of the sexually and parthenogenetically reproducing *D. mercatorum* represent the protein coding part of these genomes and that they accurately reflect chromosome organisation in this species. The gross differences in genome organisation that we see between sexually and parthenogenetically reproducing strains correspond to those typifying geographically distinct isolates [8] and do not obviously account for the ability to reproduce parthenogenetically. The sexually reproducing and parthenogenetic strains do exhibit numerous sequence polymorphisms but in the absence of any functional assay, it is impossible to know the significance of these differences with respect to the mode of reproduction.

**Fig. 2.**
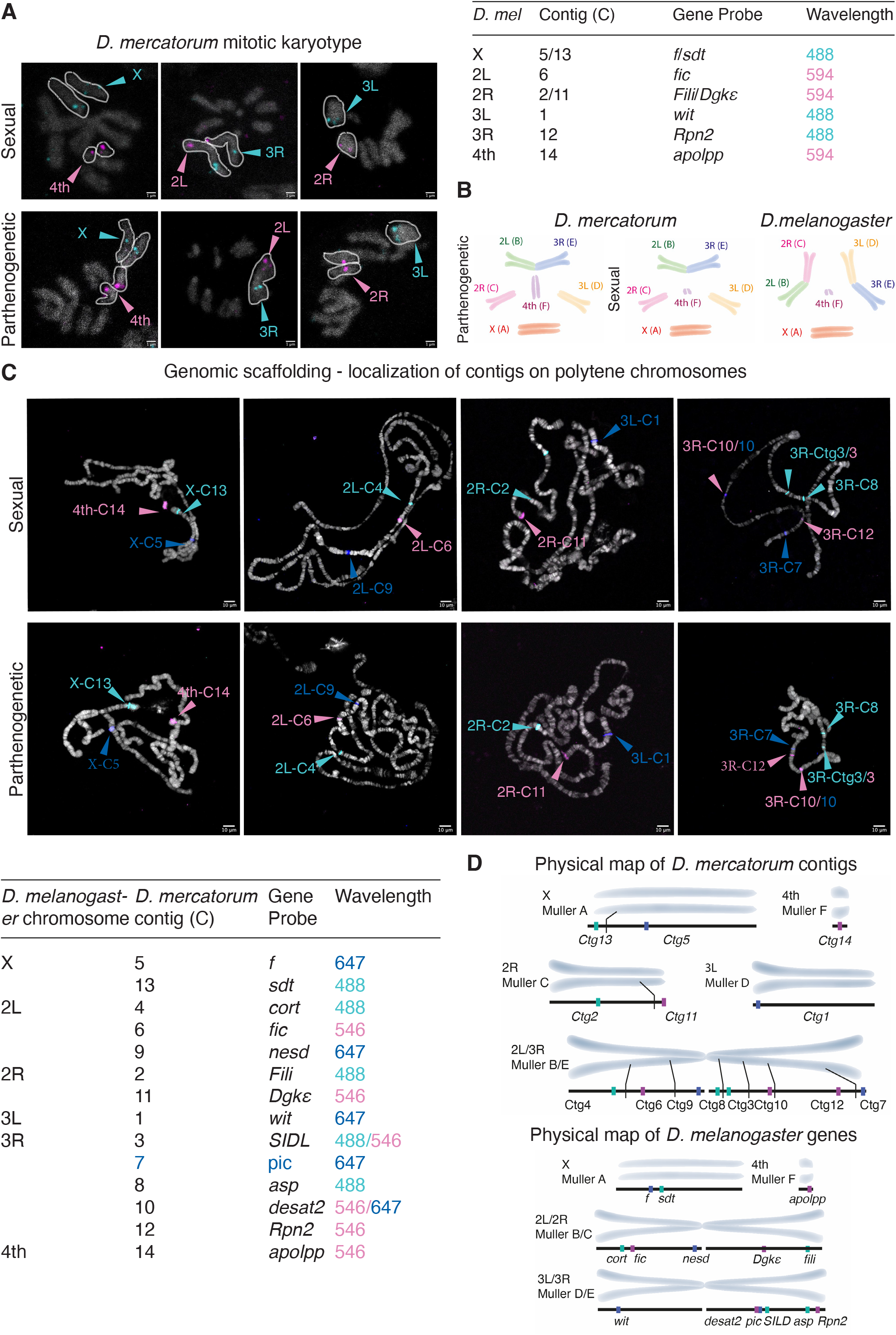
The *D. mercatorum* genome aligns with the cytological arrangement of Muller elements on mitotic and polytene salivary gland chromosomes. **(A) *In situ*** localization of the indicate gene by HCR onto mitotic chromosomes from neuroblasts of sexually reproducing and parthenogenic *D. mercatorum* larvae. Gene probes were selected to represent the indicated contigs (C). Chromosomes showing localization of the gene probe are outlined with a white line. The indicated chromosome was marked in 100% of typical karyotypes analyzed (n≥42, N≥3). The scale is 1μm. **(B)** Schematic of the arrangement of the Muller elements in sexually reproducing and parthenogenetic strains of *D. mercatorum* and in *D. melanogaster*. **(C)** Mapping of the largest 14 contigs of the sexual and parthenogenetic *D. mercatorum* genomes onto salivary gland polytene chromosomes stained with DAPI. *In situs* were carried out using the indicated HCR DNA probes corresponding to the indicated genes. The scale is 10μm. **(D)**Schematic of the mapping of the contigs onto *D. mercatorum* chromosomes and their corresponding position on *D. melanogaster* chromosomes.

### Gene expression differences between sexual and parthenogenetic D. mercatorum

We argued that genomic changes with potential to lead to changes in reproductive ability should reveal themselves as gene expression changes late in female germline development. Our knowledge of the genome then allowed us to use RNA sequencing to characterise the transcriptomes of mature eggs (Stage 14 egg chambers) isolated from the sexual, parthenogenetic, and a ‘partially parthenogenetic’ strain of *D. mercatorum*. The partially parthenogenetic strain reproduces sexually but has an enhanced ability to switch to parthenogenetic reproduction. From the three transcription profiles we identified 7656 genes that were expressed in mature eggs (Fig. S4A-B), having a similar distribution to the annotated genes (Fig. S1A-B). A comparison of gene expression profiles between the datasets identified 92 genes that were differentially expressed in all three pairwise transcriptome comparisons (Fig. S4C, Data tables S3). There were few genes that showed strong and significant differential expression in all the comparisons. A gene ontology (GO) analysis of genes with significant differential expression from all pairwise comparisons revealed enrichment of genes involved in redox, immune function, wing disc growth, biosynthesis, proteolysis, and translation (Fig. S4D). Following consideration of the GO analysis and manual curation of pairwise comparisons of gene expression, a subset was selected for further study, highlighted in the volcano plots of Figure S4E and listed in Table 2. Genes were selected that exhibited significant differential expression (*p*adj<0.05) at a level equivalent to heterozygosis (log_2_ fold change ± 0.6) or greater and which were indicated by the gene ontology analysis as being involved in common cellular processes.

**Table 2:**
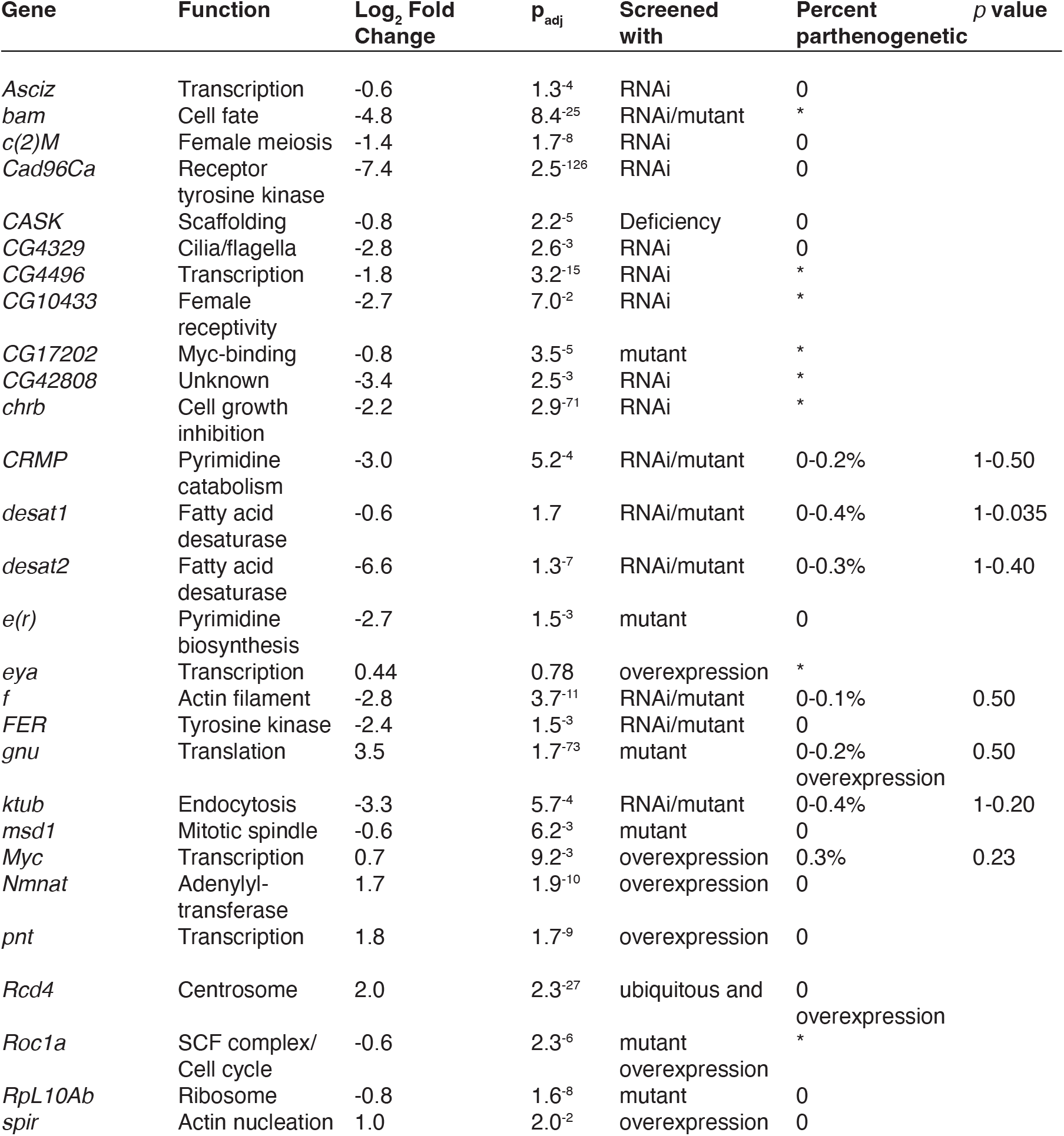
Genes differentially expressed between the parthenogenetic, partially parthenogenetic, and sexually reproducing *D. mercatorum* and the consequences of expressing their variants upon parthenogenesis in *D. melanogaster*. Gene function was assigned from flybase.org. The log2 fold change in transcript level and the *p*adj value are given only for the comparison of expression differences between parthenogenetic and sexually reproducing strains. The screen was performed with the indicated genetic tools. Results marked with an asterisk (*) are considered false positives (Supplementary information) and when there were no instances of parthenogenesis a ‘0’ is indicated. The *p* value was calculated using the Fisher’s exact test.

### Functional screens for parthenogenesis

We decided to take a two-pronged approach in an effort to identify genes that could lead to parthenogenesis in a sexually reproducing fly. The first was an unbiased screen of candidate genes identified by our transcriptomics analysis as differentially expressed between sexually reproducing and parthenogenetic strains (Table 2). The second was to screen a biased set of candidate genes. These genes were selected based on their roles in the cell and centriole duplication cycles, functions that have previously been suggested to have importance in generating diploid nuclei from the haploid products of female meiosis and in offering a maternal source of centrioles in the absence of the contribution made by the sperm basal body upon fertilization [16, 17] (Table S1). In addition to experimental controls for every gene variant screened, these screens were carried out in relation to a series of negative controls listed in Table S2. For the unbiased screen, our objective was to replicate, as far as possible, the degree of differential expression seen between *D. mercatorum* strains (Fig. S5). Since all strains of *D. mercatorum* we screened were already parthenogenetic to some degree, we carried out this screen in the non-parthenogenetic species *D. melanogaster*.Using 13 different *Drosophila* species, we first determined that a baseline indicator of parthenogenesis could be given by testing the ability of approximately 500 virgin female flies to generate progeny (Data table S4). Strong levels of parthenogenesis could be detected with as few as 30 flies. Using these criteria, we found that two typical laboratory strains of *D. melanogaster* (*W^-^* and *Oregon-R)* showed no parthenogenesis whatsoever, whereas a strain caught in the wild *(CB1)* produced a small number of embryos that showed restricted development before dying (Data table S5). This accords with previous findings that *D. melanogaster* strains caught in the wild have slight parthenogenetic ability [3].

We then tested whether down-regulating the *D. melanogaster* homologs of genes showing reduced expression in parthenogenetic *D. mercatorum* strains would result in the production of offspring that died as embryos, larvae, pupae, or from old age as adult flies. To this end, we examined CRISPR knock-out alleles that we generated in candidate genes (Fig. S6, Data table S6); publicly available mutants; or established lines in which candidate genes were down-regulated by RNAi. We also tested *D. melanogaster* constructs engineered to increase expression of genes whose homologues had elevated expression in the parthenogenetic *D. mercatorum* strains. In the case of variant alleles that were not homozygous viable, screening was carried out on heterozygotes. Together we screened a total of 47 genes (Table 2, Data table S7, and Supplementary text) and identified 16 able to cause 0.1-0.4% parthenogenesis in *D. melanogaster* (relative to number of adults screened) when their expression was either increased or decreased. The parthenogenesis observed resulted in the offspring developing to varying stages, from embryos to adult flies. In this screen in which a single gene was misexpressed, the parthenogenic offspring were largely embryos that died before hatching.

The low level of parthenogenesis detected in this single gene variant screen, where the expression of only a single gene was perturbed, is in line with earlier studies that had concluded that parthenogenesis was a polygenic trait [4]. This consideration led us to carry out a double variant screen in which we combined pairs of most combinations of variants in the 16 different genes that had been positive in the single variant screen into individual fly stocks that we then screened for parthenogenesis. This revealed several combinations able to generate between 0.5-7.4% parthenogenetic offspring that died as embryos, larvae, pupae, or from old age as adult flies (Table 3, Data table S8, and Supplementary text). From the more successful combinations, we found that one of the mutant genes either encoded a desaturase, *desat1* or *desat2*, or one of three proteins predominantly involved in regulating cell division and proliferation: *Myc, slimb*, or *polo*. Notably, 0.8% of the offspring derived from females heterozygous for a mutation in *desat2* and carrying two extra copies of a *polo* transgene expressed from its endogenous promotor (*GFP-polo*^4+^;; *desat1^-/+^)* developed to adulthood (Data table S8). This level of parthenogenesis we now observe in *D. melanogaster* is a comparable that of the ‘partially parthenogenetic’ strain of *D. mercatorum* used in generating the transcriptomics data (Data table S1).

**Table 3.** Double gene variant combinations resulting in parthenogenesis in *D. melanogaster*. Selected combinations of gene variants leading to some parthenogenetic development in *D. melanogaster* relative to two control combinations. The *p* value was calculated using the Fisher’s exact test. See supplementary information for further examples and discussion.

| Genotype | Percent<br>parthenogenetic | *p* value |
| --- | --- | --- |
| *desat1<sup>+/-</sup> / desat2<sup>+/-</sup>* | 0.4% | 1 |
| *desat2<sup>+/-</sup> / slimb<sup>+/-</sup>* | 1.2% | 0.13 |
| *Myc<sup>+/+</sup> ; desat1<sup>+/-</sup>* | 0% | 1 |
| *Myc<sup>+/+</sup> ; desat2<sup>+/-</sup>* | 0% | 1 |
| *Plk4<sup>+/-</sup> / slimb<sup>+/-</sup>* | 0.9% | 0.33 |
| *polo<sup>4+</sup> ; ; desat1<sup>+/-</sup>* | 1.7-5.2% | <0.003 |
| *polo<sup>4+</sup> ; ; desat2<sup>+/-</sup>* | 1.6-7.4% | <0.003 |
| *polo<sup>4+</sup> ; Myc<sup>+/+</sup>* | 0.6% | 0.1217 |
| *polo<sup>4+</sup> ; ; slimb<sup>+/-</sup>* | 3.1% | 3.2<sup>-08</sup> |
| *polo<sup>4+</sup> ; ; w-RNAi<sup>-/-</sup>* | 0% | 1 |
| *desat1<sup>+/+</sup> / w-RNAi<sup>-/-</sup>* | 0% | 1 |

Our results strongly suggest that parthenogenesis is associated with decreased expression of either *desat1* or *desat2*. Since the *desat2* allele is known to be a natural variant present in most populations, we determined whether our *desat1* stock carries the *desat2* allele and indeed it does. Thus, our *desat1* stock is a double mutant for *desat1* and *desat2*, accounting for its stronger phenotype.

We then asked whether the parthenogenetic offspring obtained from these screens for parthenogenesis in *D. melanogaster* were themselves able to carry out parthenogenetic reproduction and found that none of them could (Data tables S7-8). We did, however, find that the parthenogenetically produced *D. melanogaster* were still able to mate with males and produce fertile offspring (Table S3), similar to previous findings [4]. The parthenogenetically produced *D. mercatorum* offspring from the sexually reproducing stocks could also not be established as a lab stock and did not survive beyond the 7th generation of parthenogenesis, as also found previously [18]. Even our long-held stocks of fully parthenogenetic females were able to mate with males (Data table S2). Therefore, we have not found a genetic combination that leads to obligate-like parthenogenesis, but we have identified key genes for facultative parthenogenesis.

Having identified *D. mercatorum* genes whose homologues lead to a degree of parthenogenetic development when mis-expressed in *D. melanogaster*, we looked for genomic differences between the organization of these genes in sexual and parthenogenetic strains of *D. mercatorum*. We found no substantial changes in gene organisation of *desat1/2, polo*, or *slimb*, although we cannot exclude the possibility of changes in distal enhancer elements that have not been mapped (Fig. S7A-D, Supplementary text). There were several changes to the *Myc* locus that could affect the expression of the protein and change its downstream function (Fig. S7E). The Myc locus of the parthenogenetic strain showed many deletions and insertions leading to the changes in primary amino-acid sequence of the protein as indicated in Figure S8A. None of these mutations affected either the basic Helix-Loop-Helix (bHLH) DNA-binding domain, the three Myc Box (MB) domains (1-3), or the three known phosphorylation sites of the Myc protein (Fig. S7E) [19–21]. There were also changes in genome organisation at the *Myc* locus of the *D. mercatorum* parthenogenetic genome, which has a 1.4kbp repetitive region between a *Drosophila* INterspersed Elements-1 (DINE-1/INE1) transposable element (TE) and the *Myc* coding region, which are approximately 8.7kbp apart. It is possible that the 1.4kbp repetitive sequence present in the parthenogenetic genome between the gene and the TE could result in de-repression of *Myc* expression relative to the sexually reproducing flies. Finally, we detected a 48bp deletion in the parthenogenetic genome 344bp up stream of the start site which could alter *Myc* expression. The above mutations have the potential to affect the transcription, translations, or protein stability of Myc. Moreover, these mutations might also directly or indirectly perturb Myc’s functions as a transcription factor to influence the expression of the other genes identified in our study, since *Drosophila* Myc is known to broadly and subtly influence the expression of genes involved in growth, size, metabolism, apoptosis, and autophagy [22]. As *D. mercatorum* is not tractable for molecular genetic studies, future work will be required to distinguish these possibilities.

### The development of parthenogenetic embryos

To understand how development might be initiated during parthenogenesis we first examined fertilized eggs from the sexually reproducing *D. mercatorum* strain that were initiating the mitotic nuclear division cycles (Fig. S9A) for comparison to the parthenogenetic eggs. The sexually reproducing *D. mercatorum* embryos had timely nuclear divisions within the syncytium and no obvious nuclear defects. We observed that 12% of unfertilized eggs from the partially parthenogenetic strain showed one or more nuclear divisions compared to 38% of unfertilized eggs from the parthenogenetic strain (Fig 3A, Fig. S9B-G). The extent to which nuclear division cycles could take place in parthenogenetic *D. mercatorum* reflected the extent of parthenogenicity. In contrast to the sexually reproduced embryos, the unfertilized parthenogenetic and partially parthenogenetic embryos developed with abnormal numbers of nuclei (Fig. S9D) or apparent chromosomal abnormalities (Fig. S9G). All parthenogenetic *Drosophilids* appear to retain normal meiosis and rediploidise their genomes either by fusion of one or more of the four haploid nuclei arising from meiosis or by post-meiotic duplication of the haploid gamete [3, 5, 16, 23–26]. All four meiotic products, three polar bodies and the female pronucleus, are present within the *Drosophilid* egg, and the three polar bodies normally fuse and arrest in a mitotic-like state. We only observed the presence of polar bodies in 44% of parthenogenetic embryos that initiated the mitotic nuclear division cycles suggesting that the missing polar bodies may be fusing and participating in the mitotic nuclear division cycles in the developing embryos, as previously suggested [16].

**Fig. 3.**
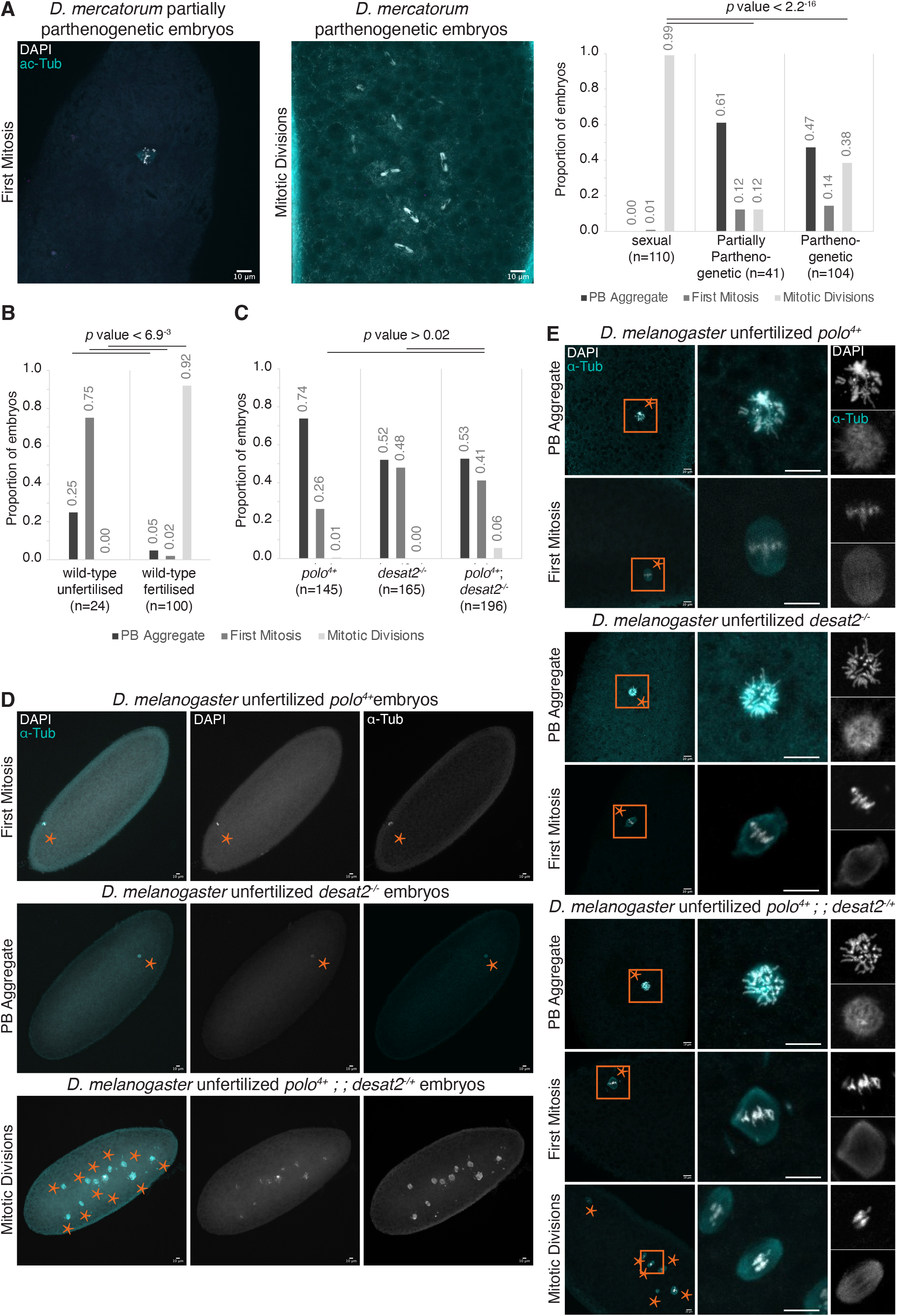
The initiation of parthenogenesis in natural *D. mercatorum* and genetically modified *D. melanogaster* embryos. (**A**) Unfertilized partially parthenogenetic and parthenogenetic *D. mercatorum* embryos showing restricted mitosis and multiple mitotic divisions, respectively. The histogram displays the proportion of sexually and parthenogenetically reproducing *D mercatorum* eggs/embryos that have polar body (PB) aggregates; initiated the first mitosis; or have undertaken multiple mitotic nuclear divisions (see also Fig. S11). (**B**) Histograms displaying the proportion of wild-type unfertilized and fertilized *D. melanogaster* eggs/embryos that have PB aggregates; initiated the first mitosis; or have entered multiple mitotic nuclear divisions (see also Fig. S12 for examples). (**C**) Histograms showing the proportion of unfertilized *GFP-polo^4+^*, *desat2^-/-^*, and *GFP-polo^4+^; desat2^-/+^ D. melanogaster* embryos that have PB aggregates; initiated the first mitosis; or have undertaken multiple mitotic nuclear divisions. (**D**) *GFP-polo^4+^*, *desat2^-/-^*, and *GFP-polo^4+^; desat2^-/+^ D. melanogaster* embryos that have initiated the first mitosis; PB aggregates; or have undertaken multiple mitotic nuclear divisions. (**E**) *GFP-polo^4+^*, *desat2^-/-^*, and *GFP-polo^4+^; desat2^-/+^ D. melanogaster* embryos that have initiated the first mitosis; PB aggregates; or have undertaken multiple mitotic nuclear divisions. Fisher’s exact test was used to calculate *p* values. Nuclei are marked with asterisks. The wild-type strain is Oregon-R. Scale bars represent 10μm.

We then examined fertilized and unfertilized wildtype *D. melanogaster* eggs and compared their development to the parthenogenetic *D. melanogaster* eggs. All unfertilized wildtype *D. melanogaster* eggs completed meiosis and 70% appeared to have entered the first mitotic division and had condensed chromosomes (Fig. S10A-B, Fig 3B). In contrast, nearly all fertilized embryos had begun to undergo the mitotic nuclear division cycles (Fig. S10C-D, Fig 3B). Development of the induced parthenogenetic *D. melanogaster* embryos mirrored the findings from the parthenogenesis screens. We focused upon embryos derived from *GFP-polo^4+^* or *desat2^-/-^* females since variants of these genes had shown the highest levels of parthenogenesis in double variant combinations. We found nearly all unfertilized eggs laid by either *GFP-polo^4+^* or *desat2^-/-^* mothers were unable to undertake mitotic nuclear division cycles (Fig. 3C-E, Fig. S11A-D). By contrast, 6% of the unfertilized eggs laid by *GFP-polo^4+^*; *desat2^-/+^ D. melanogaster* mothers could undertake at least limited mitotic cycles (Fig. 3C-E, Fig. S11E-G). These parthenogenetic *D. melanogaster* embryos were similar to the unfertilized parthenogenetic and partially parthenogenetic *D. mercatorum* embryos in that they had abnormal numbers of nuclei and/or chromosomal abnormalities. Although these parthenogenetic embryos show such abnormalities, only one normal diploid nucleus is needed to divide during early embryogenesis to generate a viable offspring. We were unable to observe any polar bodies in those unfertilized *GFP-polo^4+^*; *desat2^-/+^* - derived eggs that had initiated the mitotic cell divisions and found that number nuclei deviated from the expected 2^n^, suggesting that the polar bodies participate in the nuclear division cycles. Together, these observations lead us to propose that in these induced parthenogenetic *D. melanogaster* embryos, the recapture of polar bodies results in the diploidisation of some nuclei, which are subsequently able to capture maternally derived centrosomes, as has been proposed in *D. mercatorum* parthenogens [16, 17].

## Discussion

Our study offers the first account of a molecular basis underlying the evolution of any type of parthenogenesis in any animal. Our findings relate specifically to the Dipteran *D. mercatorum* and suggest a route through which parthenogenesis could arise in this species. By identifying genes that are differentially expressed in oocytes of sexually and parthenogenetically reproducing strains of *D. mercatorum* and modulating their expression in *D. melanogaster* while also mis-regulating nuclear and centriole/centrosome cycle genes, we have identified combinations of gene variants that enable a degree of parthenogenesis in *D. melanogaster*. Our findings suggest that an effective step towards establishing parthenogenesis is the heterozygosity of *desat1* or *desat2* coupled to overexpression of *polo* or *Myc* or heterozygosity of *slimb*, which encodes the F-box protein Slimb of the SCF ubiquitin-protein ligase.

Our results suggest a key requirement for the differential expression of *desat* and cell cycle genes between parthenogenetically and sexually reproducing strains. The ability of *desat1* mutants to enhance the phenotype of *desat2* mutants in driving parthenogenesis when heterozygous in *D. melanogaster* is likely a consequence of overlapping function between their encoded proteins, which show 85% identity in amino-acid sequence. Both *desat1* and *desat2* encode desaturases that generate double bonds in hydrocarbons and have roles in lipid metabolism. Desat1 also generates double bonds during sex pheromone biogenesis [27], and when mutated, can result in female resistance to mating. Desat2 desaturates cuticular hydrocarbons; it has been associated with increased cold tolerance [28] and is credited with imparting the ability *of D. melanogaster* to become invasive and colonise cosmopolitan habitats. The highly pleiotropic nature of *desat1* and *desat2* makes it difficult to determine how their down-regulation relates to the increased incidence of parthenogenesis. As their mutation can change phospholipid membrane composition, it is possible that this could influence a wide range of membrane associated trafficking events that could facilitate the diploidization of the haploid polar bodies, and the onset of zygotic mitoses. The potential effects of these mutations upon such events will require detailed future studies.

The other group of genes involved in enabling parthenogenesis regulate some aspect of cell cycle progression. Slimb is a subunit of the SCF, the Skp, Cullin, F-box containing ubiquitin ligase complex that controls the centriole/centrosome duplication cycle by promoting the destruction of the master regulator of centriole duplication, Polo-like kinase 4 (Plk4). *slimb* mutants are known to develop supernumerary centrosomes [29, 30]. Indeed, elevated levels of Plk4 have been shown to drive centriole duplication and function in unfertilized *D. melanogaster* eggs [31]. Such provision of maternally derived centrosomes has been shown to be a requirement for the parthenogenetic development of *D. mercatorum* embryos [16, 17].

SCF also regulates S phase entry by targeting multiple substrates. One of the multiple targets of the SCF is Myc [32]. We found only a modest increase in Myc transcripts (0.6 Log2 fold change) in the parthenogenetic strain. However, this change in expression level was highly significant (padj<0.001) and is equivalent to having one extra copy of the gene. The Myc bHLH transcription factor has been shown to give *D. melanogaster* cells a competitive growth advantage [33] and could account for the finding that parthenogenetic offspring are physically larger than the sexually reproducing animals [34]. Myc is known to promote Polo-like-kinase1 expression in mammalian cells that in turn destabilises SCF [35]. If a similar mechanism were to act in *Drosophila*, this could account for the relationships we observe here to promote parthenogenesis. Polo kinase (Plk1) plays multiple roles in the centriole duplication cycle. It is required to convert the newly duplicated centriole into a functional centrosome [36]. It also drives centrosome maturation facilitating increased nucleation of microtubules during mitosis [37, 38]. Thus, we anticipate that a doubling of Polo kinase levels in the *Drosophila* egg will facilitate the formation and mitotic function of centrosomes that are required for parthenogenesis.

Although we have shown that we can induce parthenogenesis in a sexually reproducing line of *D. melanogaster* to a similar degree as a partially parthenogenetic strain of *D. mercatorum*, we are not able to maintain these animals as a parthenogenetic stock. Therefore, although we identify a significant step towards heritable parthenogenesis, this is not the end of the story and additional changes would be required for parthenogenesis to become fixed in a population and transit to more obligate-like parthenogenetic reproduction. Moreover, there are likely to be many alternative paths to the devolution of sexual reproduction in animals and this could explain the varying degree of parthenogenetic ability, not only within *D. mercatorum*, but also across the *Drosophila* genus. Given the polygenic nature of facultative parthenogenesis and the fact that there are multiple inputs into core cell cycle regulation, it may explain why no unifying signature of parthenogenesis has been found to date [39]. Thus, we anticipate that parthenogenesis might have different causal events in each species or even between individuals of the same species.

Some consider sporadic facultative parthenogenesis to be an unimportant accident. However, there could be a benefit of having sporadic facultative parthenogenesis inducing heterozygotic mutations floating around in the population, they may facilitate an ‘extinction escape hatch’. Such a possibility could help a lineage of the species stave off extinction in the face of isolation until an opportunity to mate arises again. Parthenogenesis is spread across the order Diptera and rare facultative parthenogenesis is prevalent in *Drosophila* [40] making it likely that the mechanism we propose is not restricted to *D. mercatorum*.

## Supporting information

DataTableS1

DataTableS2

DataTableS3

DataTableS4

DataTableS5

DataTableS6

DataTableS7

DataTableS8

## Acknowledgements

We would like to thank Isabel Palacios, John Welch, and Sally Otto for reading the manuscript and scientific discussions. We would like to thank Richard Durbin for advice, discussions, and computation assistance and Frank Jiggens for advice and discussions. We would like to thank Jonathan Day (Frank Jiggens’ Lab) and Bettina Fischer (Richard Durbin’s Lab) for advice on library preparation, technical assistance, protocols, reagents, and endless patience. We would like to thank Harry Choi and Mike Liu, from Molecular Instruments for troubleshooting and for gifted reagents that were during the optimisation of the HCR FISH protocol for use on DNA. We would like to thank Paula Almeida-Coelho for technical advice on in situs and karyotyping. We would like to thank Frank Sprenger for flies, kindly brought to Cambridge. Finally, we would like to thank the Genetics Fly Facility for embryo injections, particularly Dr Alla Madich for multiple attempts at developing transgenics in *D. mercatorum*.

## Data and materials availability

All raw and analyzed *D. mercatorum* genomic and transcriptomic data generated by this study will be provided on public databases, all code used for analysis in on GitHub via FabianDK and ekg, all *Drosophila strains/stocks* will be offered to Bloomington Stock Centre or made available upon request.

