## SupplementaryMaterials for "Virgin Birth: A genetic basis for facultative parthenogenesis"

**Materials and Methods**

*Drosophila* Stocks

All stocks that were screened have their origin or stock number given in the tables with the data (Data table S1, S4-S6). The stocks used for CRISPR generated alleles, complementation test, and rescue are listed below.

*Drosophila* CRISPR

The CRISPR/Cas9 stocks were created using *nos-cas9* and *act-cas9* from Fillip Port crossed with a transgenic line expressing the gRNA line [1]. The CRISPR Optimal Target Finder (http://tools.flycrispr.molbio.wisc.edu/targetFinder/) was used to design the 20mer target sequence. The transgenic gRNA flies were created with either $y^1$ $sc^1$ $v^1$ *P{nos-phiC31\int.NLS}X; P{CaryP}attP2* (BDSC 25710) or $y^1$ $v^1$ *P{nos-phiC31\int.NLS}X; P{CaryP}attP40* (BDSC 25709). The gRNA sequences are given in Data table S6. Two gRNAs for each gene were transformed into pCFD4 from Fillip Port [1]. The plasmid containing the gRNA sequence were injected into embryos by the Genetics Fly Facility, University of Cambridge, thereby creating the transgenic gRNA-expressing gRNA fly stocks. Flies from these stocks were crossed to *cas9* expressing females and after two rounds of 'balancing' and mutant screening, the mutant stocks were established. To screen for mutants, genomic DNA was isolated from flies stocks by crushing 10 flies in 200 µL of BufferA (100 mM Tris pH 9.0, 100mM EDTA containing 1% SDS), and incubating the lysate at 65-70°C for 30 min. The lysate was chilled on ice for 5 min before adding 30 µl of 7.5M $NH_4$-acetate, mixed, and chilled on ice for 20 min. The lysate was then centrifuged for 15 min at 13,000 rpm. The pellet was removed with a tip, 150 µl of isopropanol added, and centrifuged for 10 min at 13,000 rpm. The supernatant was decanted, and the DNA pellet was washed with 70% ethanol. The DNA was then dissolved in 100 µl of $dH_2O$. 1 µl was used per PCR reaction.

PCR was performed with the primers listed in Data table S6. PCR products were run on a 1% agarose gel and mutants were sent for sequencing.

The FRT chromosome stocks use were P{ry[+t7.2]=neoFRT}19A; ry[506] (BDSC 1709) or *w*; P{w*$^{+mW.hs}$*=FRT(w*$^{hs}$*)}2A* (BDSC 1997).

The stocks used for complementation were *asl* deficiency: *w1118; Df(3R)ED5177, P{3'.RS5+3.3'}ED5177/TM6C, cu1 Sb1* (BDSC 8103), morula mutants: *px1 bw1 mr1 sp1/In(2LR)bwV1, ds33k bwV1* (BDSC 380) and *chl1 l(2)bw1 bw2b mr2/SM5* (BDSC 257), *Plk4* deficiency: *w[1118]; Df(3L)BSC418/TM6C, Sb[1] cu[1]* (BDSC 24922), *plp* deficiency: *w[1118]; Df(3L)BSC837/TM6C, Sb[1] cu[1]* (BDSC 27916), *Sas-6* deficiency: *w[1118]; Df(3R)BSC794, P+PBac{w[+mC]=XP3.WH3}BSC794/TM6C, Sb[1] cu[1]]* (BDSC 27366), slimb deficiency: *w[1118]; Df(3R)BSC508/TM6C, Sb[1] cu[1]* (BDSC 27916).

The stocks used for rescue experiments were: *ana2* rescue: *pUbq-ana2 / CyO* (Glover Lab), asl rescue: *UASp-Asl-WT/CyO* (Glover Lab), *Plk4* rescue: *w; UASp-Sak/Cyo ; MKRS/TM6B* (Glover Lab), *plp* rescue: *Ub-plp::GFP5.2/CyO; TM3, sb/TM6B* (Ruslan Lab), *sas-6* rescue: *pUbq-sas-6 / CyO* (Glover Lab).

Parthenogenesis Assay

The test for parthenogenesis was adapted from Stalker 1954 [2]. Batches of 1-70 virgins were collected and maintained for the duration of their lives on fresh food that was changed weekly. The old food tube was kept for more than 3 days and then examined for parthenogenetic development by the presence of brown embryos or further development. All screening for offspring was carried out blind, where all sample labels were kept with the flies on tape with their fresh food and the old food tubes were mirrored on a second tray for >3 days prior to screening. If there was evidence of further development, then the tube was kept, and the animal was left to reach its final stage of development. If a parthenogenetic fly was produced, it was then maintained for the duration of its life on fresh food. We documented how many instances of parthenogenesis occurred relative to the number of adults screened since the number relative to eggs laid were extremely small and harder to conceptually rectify. For example, 1 fly per 100,000 eggs laid is a small number, in contrast 1 fly per 100 adult flies screened gives a greater impression about the prevalence and potential to populate an area.

Wolbachia Test

PCRs were run against *D. mercatorum* genome preps using general primers (wsp 81F: TGGTCCAATAAGTGATGAAGAAAC, wsp 691R: AAAAATTAAACGCTACTCCA) against Wolbachia and the resulting DNA fragments were subjected to electrophoresis on a 1% agarose gel. The primer sequences and positive control were provided by Julien Martinez, Frank Jiggens Lab, Cambridge.

*desat2* mutation Test

PCRs were run against genome preps for the *desat1/2 D. melanogaster* stocks used for screening parthenogenesis using primers to the 5'UTR of *desat2* (desat2-UTR-FWD: AAGAGCTCGCCAGCTATCTAC, desat2-UTR-REV: AAGGACACCCGTTTCTCTGG). The resulting DNA fragments were subject to electrophoresis on a 1% agarose gel, purified, and sequenced and then compared to *D. melanogaster* (assembly Release 6 plus ISO1 MT), which does not have the 16bp deletion 150bp upstream of the start codon.

Hybridisation Experiments

The reciprocal crosses of males and females from different *D. mercatorum* strains were maintained for the duration of their lives with their food changed weekly. If offspring (F1) were produced, they were checked for the presence of at least 3 males, since *D. mercatorum* does produce non-disjunction males (X/0) by parthenogenesis [3]. The F1 were then flipped into a new tube and checked for male and female offspring (F2). If F2 were produced, then both the males and the females were fertile.

Short- and Long-read Genomic Library Preparation

For the *D. mercatorum* genomes, long-read sequencing data were used to generate scaffolds and Illumina data were used to polish the assembly.

*Long-read Nanopore Library Preparation:*

Parthenogenetic Genome: The parthenogenetic isofemale *D. mercatorum* line 15082–1525.07 was obtained from the National Drosophila Species Stock Centre (Cornell University). A single female was isolated to set up a new isofemale line that was then amplified. High molecular weight DNA was extracted from 32 females from the new isofemale line. To prepare high molecular weight DNA, flies were homogenized in a 1.5 ml microfuge tube containing 150 μl of SDS buffer using a pestle driven by a handheld electric mixer for 10 seconds, 350 μl of SDS buffer (0.5% (w/v) SDS, 0.200 M Tris, 0.25 M EDTA, 0.250 M NaCl) was added, and the mixture was incubated at 37°C for 4 h. 5 μl of RNase A (100 mg/ml) was added, and the mixture was incubated at 37°C for an additional 2 h. 5 μl of Proteinase K (20 mg/ml) was added, and the solution was incubated for a further 2 h at 50°C. The DNA was then extracted using 240 μl of the phenol layer of phenol/chloroform/isoamyl alcohol (25:24:1), stabilized, saturated with 100 mM Tris-EDTA to pH 8.0, Acros Organics, from Fisher Scientific. Following mixing on a gentle agitator at room temperature for 3 min, the mixture was centrifuged at 12,000 x g for 10 min, and the supernatant poured into a new 1.5 ml tube. The phenol extraction was repeated once more followed by a final chloroform extraction. The DNA was precipitated by adding 500 µl -20ºC absolute ethanol, incubating at -20ºC for 5 min, before removing the DNA precipitate. The DNA was placed into a new tube with 70% ethanol and centrifuged at 12,000 g for 3 min and the pellet washed again with 500 ml of 70% ethanol before centrifugation at 12,000 g for 3 min to remove residual salt. The pellet was then dried at 37ºC for 30 min and 50 µl of elution buffer (10mM Tris, pH 8.0, in Nuclease free water) was added. The DNA was left at room temperature for 3 days in order to enable dissolution prior to sequencing. 457.2 ng of isolated DNA was then sequenced on the Nanopore MinIOn using the SQK LSK-109 ligation protocol and FLO-MIN106D R9 Version Spot-ON Flow Cell RevD.

Sexually Reproducing Genome: The sexual *Drosophila mercatorum* line 15082–1511.00 was obtained from the National Drosophila Species Stock Centre (Cornell University). High molecular weight DNA was extracted from 32 adult females. A slightly modified QIAGEN Genomic-tip (100/G) protocol was used for library preparation. Briefly, we prepared lysis buffer (9.5 mL Buffer G2 and add 19 µL Qiagen RNaseA, mix), added 32 *D. mercatorum* flies directly into 1.5ml tube, place in liquid Nitrogen ($N_2$), and homogenised with a pre-cooled pestle to a fine powder. Samples were kept cold with liquid Nitrogen or dry ice the entire time. 1ml of lysis buffer was added to the 1.5ml tube and then transferred a to 15 mL tube (on ice), repeated 2x and then the remaining lysis buffer was added to the tube. We immediately vortexed for 5 seconds and then incubate at 37°C for 1 h, gently inverting tube after 30 min. We added 500 µL Qiagen Proteinase K, gently invert tube, and incubated at 50°C for 2 hours, gently invert tube after 1 h. 10 minutes before the lysis incubation time ends, we prepared the genomic-tip by equilibrating it with 4 mL Buffer QBT. We vortexed the lysed sample for 5-10 sec at max speed and then gently transferred the sample into 5x 2 mL tubes by pouring. The samples were then centrifuge at 5,000 g for 10 min at 4°C. Immediately after the centrifugation the samples were poured on the equilibrated genomic-tip. We washed with 2 x 7.5 mL Buffer QC. We pre-warmed 5 mL Buffer QF to 50°C and eluted the gDNA with 5 mL warm Buffer QF into a 50 mL tube. 3500 µL Isopropanol was added to the sample and gently invert 10-20x. We gently divide the sample into 2.0 mL tubes by pouring and centrifuged immediately at 10,000g for 30 min at 4°C. We washed the pellet 2x with 1 mL cold 70% Ethanol and spun at 10,000g for 5 min at 4°C. The DNA was air dried for 5-10 min, re-suspended in 50 µL EB, incubated at 37°C for 30-60 min, and then left in the fridge overnight. We very gently flick-mix the tubes and quantified the sample using Qubit and Nanodrop and ran on the Agilent Tapestation. 2.57 µg of isolated DNA was then sequenced on the Nanopore MinIOn using the SQK LSK-110 ligation protocol and FLO-MIN106D R9 Version Spot-ON Flow Cell.

*Short-read Illumina Library Preparation:*

Parthenogenetic Genome: Illumina short-read whole genome sequence data was generated for the parthenogenetic female *Drosophila mercatorum*. DNA was extracted from a single female from the same isofemale line used for the nanopore DNA sequencing. The fly was homogenized in a 1.5 ml microfuge tube containing 150 µl of lysis buffer using a pestle driven by a handheld electric mixer for 20 seconds; 350 µl of SDS buffer was added and the mixture incubated at 37°C for 2 h. 5 µl of RNase A (100 mg/ml) was added and the mixture incubated at 37°C for an additional 2 h. 5 µl of Proteinase K (20 mg/ml) was then added and the solution incubated at 50°C for an additional 1 h. The DNA was then extracted with 240 µl of the phenol layer from phenol/chloroform/isoamyl alcohol (25:24:1), stabilized, saturated with 100 mM Tris-EDTA to pH 8.0, Acros Organics, from Fisher Scientific. Following mixing by hand at room temperature for 1 min, the mixture was centrifuged at 12,000g for 10 min, and the supernatant poured into a new 1.5 ml tube. The phenol extraction was repeated once more, followed by extraction using the chloroform layer. The DNA was precipitated by adding 500 µl of -20°C absolute ethanol. Following centrifugation at 12,000g for 5 min, the pellet was washed twice in 70% ethanol and then dried at 37°C for 30 min before addition of 52.5µl of elution buffer (10mM Tris, PH 8.0, in Nuclease free water). 52.5 µl of DNA was put into a Covaris Screw Cap microTUBE for M220 and sonication performed on an E220evolution Focused-ultrasonicator. The library was then prepared using the KAPA HyperPrep Kits for NGS DNA Library Prep by Roche and sequenced on a NovaSeq with 150 bp PE reads.

Sexually Reproducing Genome: Illumina short-read whole genome sequence data was generated for the sexual female *Drosophila mercatorum*. The same protocol was used as for the parthenogenetic short-read DNA preparation. However, the library was then prepared using the NEBNext® Ultra™ II DNA Library Prep Kit for Illumina® by New England Biolabs and sequenced on a MiSeq with 150 bp PE reads.

Genome Assembly

The genomes for the wildtype and parthenogenic strains were assembled using wtdbg2 [4] version 2.5 using the setting “wtdbg2 -x ont -g 200m” and following the consensus generation procedure (wtpoa-cns, minimap2 [5], samtools [6]) recommended by the author. We then followed a polishing procedure similar to that used in the Vertebrate Genomes Project (VGP), wherein alignments of Illumina reads were compared to the genome assemblies and used as input to freebayes [7]. bcftools consensus [8] was used to correct the genome to match homozygous calls from the Illumina alignments. The assemblies have N50s of 22.7 Mbp for the wildtype and 16.4 Mbp for the parthenogenic strain.

Alignment

The alignment of the nanopore reads back to the generated assemblies was used for evaluation of assembly quality. We completed the alignment using minimap2 with settings "-a -x ont", resulting in alignments for each input nanopore read set to its respective assembly. As a basic evaluation of assembly quality, we compared the two *D. mercatorum* assemblies to each other using whole genome alignment with wfmash. The resulting alignment covers 151.14Mbp, including 142.25Mbp of matching base pairs. The gap compressed identity (often quoted to describe species level divergence) is 98.67%, while the "block" identity is 94.11%. This suggests that the base-level accuracy of each assembly must be significantly less than around 1%, as we expect evolutionary divergence of around that seen in the gap compressed identity. We additionally completed whole genome alignments with wfmash [9] to evaluate the relationship between the assemblies and with the *D. melanogaster* reference assembly, using "wfmash -p 70" to obtain alignments at up to 30% divergence. The gap compressed identity metric produced by wfmash is 75.85% (over 75.28Mbp) between the wildtype *D. mercatorum* and *D. melanogaster* assemblies and 75.55% (over 73.44Mbp) between the parthenogenic *D. mercatorum* strain and the *D. melanogaster* reference genome. The high level of rearrangement in this lineage, and ~47Mya divergence results in a highly fragmented alignment. However, for both whole genome alignments between the *D. mercatorum* strains and *D. melanogaster* reference, chromosome level gene content appears to be maintained, despite a very high level of rearrangement.

Coverage

Evaluation of the coverage of alignments was completed using bedtools [10]. Windows for each 10kb of the assembly were established using "bedtools makewindows", and in each we computed the coverage of alignment using “bedtools coverage -mean”. We find the median coverage of a 10kb window to be 87.73x in the wildtype assembly and 95.74x in the parthenogenic assembly. As shown in plots the coverage is even across all large contigs in the assemblies, which comprise the vast majority of the assembled sequence, and most deviations in coverage occur in shorter contigs that may represent possible misassemblies or collapsed repeats. We do find that the wildtype assembly contains some contigs at lower than expected coverage, possibly corresponding to heterozygosity in this non-inbred strain.

Nucmer and gene distribution

As a complementary approach to the dot plots, we further used nucmer (MUMmer v3.23) [11]with options --coords --maxgap 500 –maxmatch to align contigs from *D. mercatorum* to *D. melanogaster* chromosomes (reference version v6.27). To avoid potential misalignments caused by repeats, both genomes were hard-masked using RepeatMasker [12] (see Annotation below) before running nucmer. The show-coords command with -l -c options was then used to create a coordinates file. We considered that a *D. mercatorum* contig maps to a *D. melanogaster* chromosome if >40% of the alignments map to one *D. melanogaster* chromosome, while all other chromosomes have <20% alignments.

Genome Annotation

To annotate the *D. mercatorum* genome assembly, we first identified repeats in the genome assembly using RepeatModeler2 (v 2.0.1) [13] with option -LTRStruct. We then soft-masked repeats in the genome utilizing the found repeats using RepeatMasker v4.0.9 [12] with options -gccalc -s -nolow -norna -gff -xsmall (or without -xsmall for hard-masked assembly used in nucmer analysis, see above). Next, we trimmed RNA-seq paired-end reads (see Transcriptomics below) using cutadapt v2.0 [14] and options -a AGATCGGAAGAGC -A

AGATCGGAAGAGC -q 20 --minimum-length 50. Trimmed reads were mapped against the unmasked genome assembly using STAR v2.7.0e [15] and bam files from parthenogenic/partial parthenogenic/sexual lines merged using samtools *merge* [6]. To predict genes, we employed BRAKER2 [16] using the soft-masked genome assembly (with option --softmasking) and the merged bam file. To identify most likely 1to1 homologs between *D. mercatorum* and *D. melanogaster* genes, we first extracted the CDS sequences from the identified *D. mercatorum* genes using gffread and obtained the protein gi numbers from *D. melanogaster* with esearch and efetch (July 2020). We then used BLASTx (ncbi-blast-2.10.1) to blast the *D. mercatorum* CDS against the *D. melanogaster* proteins with options -db nr -num_threads 60 -outfmt '6 qseqid stitle qlen slen length qcovs qcovhsp evalue bitscore pident sacc sseqid sscinames' -max_target_seqs 137871. The blast output was filtered for entries with an evalue of <1e-10, minimum query coverage >50 and percent identity >35. Gene names and FBgn numbers were added to accession numbers utilizing mappings obtained from FlyBase (fbgn_NAseq_Uniprot_fb_2020_03.tsv). To get reverse hits, we used tBLASTx to blast the *D. melanogaster* CDS (reference version 6.35) against the *D. mercatorum* CDS with the same options as above except from -max_target_seqs 18313 and filtered the output using the same cut-offs.

Mitotic chromosome preparation

Brains were dissected from 3rd instar larvae in saline (0.7%NaCl) and then subjected to hypotonic shock by incubation in 0.5% trisodium citrate for 9 min. The brains were then fixed by a 60 sec incubation in 45% acetic acid followed by 5 min in 60% acetic acid on a coverslip. A slide was placed over the coverslip and squashed between two sheets of blotting paper. Immediately after squashing, the preparation was frozen in liquid $N_2$ and the coverslip removed using a scalpel. The squashed brain preparation was then dehydrated by successive

5 min incubations in 70% and 100% ethanol and then air-dried. Prior to denaturation, slides were baked at 58°C for 1 h in a dry oven. The preparations were then denatured with 70% formamide in 2×SSC at 70°C for 20 min and subjected to further dehydration by successive 5 min incubations in 70% and 100% ethanol, air-drying and immediate application of the HRC protocol.

<u>Polytene chromosome preparation</u>

Salivary glands were dissected from 3rd instar larvae in saline (0.7%NaCl) and then fixed in 45% acetic acid for 30-60 sec. Glands were transferred to a 6 µl drop of 1:2:3 lactic acid:water:acetic acid on a 18mm square cover slip, which was covered with a slide and squashed. Immediately after squashing, the preparation was frozen in liquid $N_2$. The coverslip was removed with a scalpel, the preparation dehydrated by emersion three times for 10 min in 95% ethanol before finally being air-dried. Chromosomes were denatured by incubating the slides in 2x SSC at 65°C for 30 min and then twice in 2 x SSC at room temperature for 10 min before incubation in 70 mM NaOH for 3 min. Slides were rinsed in 2x SSC and then soaked twice in 70% EtOH for 5 min, twice in 95% EtOH for 5 min, and then air dried before immediately preceding to the HRC protocol.

<u>Molecular Instruments HCR Protocol</u>

All materials including buffers and probes were purchased or gifted from Molecular Instruments. To ensure optimal hybridization to mitotic chromosome preparations, probes were selected for accessible gene regions by the criteria that the chosen genes were highly transcribed in brain tissue. For multiplexed HCR, samples were pre-hybridized by incubating with 200 µL of probe hybridization buffer for 10 min at 37°C inside a sealed plastic box with damp paper towel. The probe solution was prepared by adding 0.4 pmol of each probe set (by taking 0.4 µL of 1 µM stock) to 100 µL of probe hybridization buffer pre-warm 37°C. The pre-hybridization solution was removed and excess buffer drained form the slide by blotting its edges on a Kimwipe. 100 µL of the probe solution was added to the top of the sample, which was covered by a coverslip for overnight incubation for 20 h in the 37°C in a sealed plastic box with damp paper towel. The slide was immersed in probe wash buffer at 37°C to float off coverslip and excess probe removed by incubating the slide at 37°C sequentially in: (i) 75% of probe wash buffer : 25% 5x SSCT for 15 min (ii) 50% of probe wash buffer : 50% 5x SSCT for 15 min (iii) 25% of probe wash buffer : 75% 5x SSCT for 15 min (iv) 100% 5x SSCT for 15 min. The slide was then immersed in 5x SSCT for 5 min at room temperature before drying by blotting its edges on a Kimwipe. 200 µL of amplification buffer were added onto the top of the sample, which was then "pre-amplified" in a sealed plastic box with damp paper towel for 30 min at room temperature. 6 pmol of hairpin h1 and 6 pmol of hairpin h2 were prepared by snap cooling 2 µL of a 3 µM stock that was pre-heated at 95°C for 90 seconds and cooled to room temperature in a dark drawer for 30 min. The hairpin solution was prepared by adding snap-cooled h1 hairpins and snap-cooled h2 hairpins to 100 µL of amplification buffer at room temperature. The pre-amplification solution was removed and excess buffer drained from the slide by blotting its edges on a Kimwipe. 100 µL of the hairpin solution was added to the top of the sample, which was covered with a coverslip sample and incubated for 20h in a sealed plastic box with damp paper towel at room temperature. The slide was immersed in 5x SSCT at room temperature to float off the coverslip and excess hairpins removed by incubating the slide in 5x SSCT at room temperature in one 5min; two 30min; and two 5min washes. The slide was dried by blotting its edges on a Kimwipe. 2 drops of Vactashield containing DAPI was added to cover the specimen, which was then covered with a coverslip and sealed with nail varnish.

RNA Library Preparation

All libraries were prepared together and there were 3 biological replicates. To characterize gene expression in *D. mercatorum*, batch transcriptional profiling was performed upon Stage 14 egg chambers dissected from virgin female flies that were evenly distributed between 3-12 days old. This age range was chosen because the parthenogenetic females have their maximum output of eggs that hatch when they are 7-14 days old (week 2), whereas the partially parthenogenetic tend to lay eggs that hatch with a bimodal distribution with maxima in week 1 and week 3 (data not shown and Data Table S1). All animals for each sample were kept together and treated identically prior to dissection. The mothers of the flies that were dissected were all approximately the same age and the dissected animals were all well fed and controlled for population size while they were developing. The virgins were stored at 25°C after collection. Stage 14 egg chambers were dissected from all flies on the same day and in batches controlling for time of day. The ovaries were placed in a new well and 4 stage 14 egg chambers were taken from the ovaries of each fly for each sample in batches of 10 flies at a time. The dissection was carried out in 0.1% PBST and the egg chambers were individually transferred to a 1.5ml tube containing ice cold 0.1% PBST upon isolation. Once the target amount was reached, PBST was removed and RNALater solution (200 µl) was placed into the tube to fix the egg chambers and stabilise the RNA. The tubes were stored at 4°C overnight. The RNALater was removed, and the RNA was prepared using the standard protocol of the RNAEasy kit from Qiagen. Libraries were prepared using the standard protocol of the KAPA mRNA HyperPrep Kit from Roche. The RNA libraries were sequenced on a NovaSeq with 150 bp PE reads.

Transcriptomics

To detect changes in expression, transcriptional profiling was carried out in triplicate for three different conditions: parthenogenetic *D. mercatorum*; sexually reproducing *D. mercatorum* that naturally have a high frequency of parthenogenesis; and sexually reproducing *D. mercatorum* that do not have a naturally high frequency of parthenogenesis. We used STAR v2.7.0e to map trimmed, paired-end reads against the *D. mercatorum* assembly employing the gene annotations (gtf file) created by BRAKER2 (see above) and counted reads mapping to exons using featureCounts (subread v1.6.3). We filtered out genes that had 0 counts across all samples, and then used DESeq2 fitting 'parthenogenesis condition' as factor and contrasting pairwise expression differences across the three conditions. P values were corrected for multiple testing using FDR.

Gene Ontology (GO) analysis

We performed GO enrichment (Fisher's exact test) and gene set enrichment analysis (GSEA, Kolmogorov-Smirnov test) using the weight01 algorithm in topGO [17]. As there are no GO annotations available for *D. mercatorum*, we employed the corresponding *D. melanogaster* homologs or orthologs (see above) and therefore assumed that genes would have a similar function and protein sequence in the two species. For *D. mercatorum* genes that mapped to multiple *D. melanogaster* genes, we randomly picked one *D. melanogaster* gene, and accounted for differences in enrichment by performing a total of 100 GO analysis iterations. For each iteration, we considered GO terms significant at an FDR (false discovery rate) <0.1 and report the frequency depicting how often a GO category fell below this FDR threshold. The analysis was performed for all pairwise parthenogenesis-type comparisons, considering either all or only up or downregulated genes.

Embryo Preparation

Mothers were aged for at least 3 weeks prior to initiating embryo collections. Unfertilized embryos were collected for 2 h at 25°C and then incubated at 25°C for 2 h. Fertilized embryos were collected for 2 h and immediately fixed. The embryos were dechorionated in 50% bleach for 3min and then washed. The embryos were then transferred to either a 50:50 100% heptane:4% paraformaldehyde/PBT mixture or a 50:50 100% heptane:100% methanol mixture and fixed for 20 min on a rotator. Then the bottom layer was removed, methane added to the remaining liquid, and then then all the liquid was removed. The embryos were washed 3 times with 100% Methanol and then slowly and gradually rehydrated by successive immersions in 25%, 50%, 75%, and 100% PBS. The rehydrated embryos were then washed 2x with PBS containing1%Tween (PBT), blocked by incubating for 1 h in PBT containing 10% BSA before incubation with the primary antibody in PBS containing 2% Tween and 1% BSA for 16-24 h at 4°C. After washing the embryos three times with PBT for a total of 30 min, embryos were incubated with the secondary antibody for 4 h at room temperature, or 16 h at 4°C. Finally, the embryos were washed three times with PBT for a total of 30 min and mounted in Vectashield (Vector) containing DAPI to visualize DNA.

**Primary antibodies:** mouse α-acetylated-tubulin, clone 6-11B-1 (1:200), mouse α-tubulin, DM1A (1:200), rabbit α-Histone H2A antibody, ab13923 (1:500) **Secondary antibodies:** (all 1:500) Goat α-Mouse 488 and 647 from Life Technologies, Goat α-Rabbit 488 from Invitrogen and 647 from Life Technologies.

Imaging

All images were acquired on a Leica SP8 confocal microscope, and the images were minimally optimized for brightness and contrast using ImageJ. No other image alteration was performed. Nearly all images presented are projections of multiple focal planes.

Statistical Analysis

The statistical analysis used for the functional screens was made using the Fisher's Exact Test. This test was selected because it is permissive to having samples with '0' instances of positive cases in the controls. For the single gene variant screen, the control used was RNAi stock, the MTB (III) gal4 stock, the *w* mutant, or the Ubi-Rcd4 stock. The last 3 stocks showed no instances of parthenogenesis. For the double gene variant screen, the control is either given in the table or it was the strongest single gene variant causing parthenogenesis in the given combination.

### Supplementary Text

The parthenogenesis screen detailed analysis

i) Overview

We performed two screens of genes identified as showing differential expression from the *D. mercatorum* transcriptomic studies and from a curated list of cell cycle/centrosome genes for their ability to cause parthenogenesis when their expression level was increased or decreased in *D. melanogaster*. For these screens we tracked the temperature at which flies were maintained, the number of crosses if applicable, the number of flies collected, the maximum lifespan per batch, the average maternal age of parthenogenesis, proportion of life completed at the age of parthenogenesis, the maximum developmental stage reached by parthenogenetic offspring, proportion of offspring produced of the overall number of flies screened, and any additional observations about any adult flies generated, and *p* value when appropriate.
We screened existing mutant alleles for the selected genes when homozygous; newly created CRISPR alleles, knockdowns resulting from RNAi using existing RNAi fly lines; and flies carrying over-expression constructs. In our experience, RNAi is not completely reliable in the female germline even in the VALUM20/22 collection designed for germline RNAi. We also

cannot exclude the possibility that differential expression we tracked in *D. mercatorum* could be more widespread than in just the germline, in which case driving RNAi in the *D. melanogaster* germline alone might be insufficient to achieve the parthenogenesis phenotype, since the mother contributes to the egg directly from the fat body and the soma also contributes to the germline. Notwithstanding these non-sequiturs, we will discuss the outcome of RNAi screens below.

ii) Controls

The following Gal4 drivers for overexpression or RNAi were tested as controls for parthenogenesis: *bag of marbles (bam) gal4*; *maternal tubulin (MTB) gal4* on the 2$^{nd}$ and 3$^{rd}$ chromosomes; and *nanos (nos) gal4.* Both *MTB-gal4* and *nos-gal4* stocks showed a small degree of parthenogenesis. We also screened the double balancer stock that we used to balance any stocks that we used for our double variant screen. It too showed a low level of parthenogenesis.

Five negative controls were used (Table S2): 1) a mutant for CG3436, a gene that was expressed in the egg and implicated in the cell cycle but not differentially expressed, 2) two mutant alleles for *dhd*, another gene that is necessary for the onset of embryogenesis, 3) a mutant and RNAi line for *Klp64D*, a gene that is not involved in the cell cycle and not differentially expressed, 4) a mutant/RNAi line for *Trx-2*, a gene that is necessary for the onset of embryogenesis, and 5) a mutant and RNAi for *white* (*w*), which was not expressed in the *D. mercatorum eggs*. A TE insertion allele for CG3436, showed no parthenogenesis as expected. The null and p-element alleles for *dhd* also did not show any degree of parthenogenesis. A hypomorphic allele for *Klp64D*, did not show any degree of parthenogenesis, whereas the *Klp63D* RNAi line showed a little parthenogenesis when homozygous when the RNAi stock was homozygous (not combined with the gal4) but there was no parthenogenesis when its expression was driven by Gal4 and is therefore considered a false positive (see below). The fourth negative controls comprised 2 RNAi lines and a mutant for *Trx-2*. One *Trx-2* RNAi line did not show any degree of parthenogenesis and the other showed a low level of parthenogenesis when expression of the RNAi was driven by Gal4. This was considered an ambiguous result. However, since a TE insertion allele did not show any degree of parthenogenesis, the positive RNAi line could be the consequences of an off-target effect. This gene was flagged for further testing in the secondary screen as a precautionary measure. Finally, a classic null $w^{-/-}$ mutant that was also included as part of the preliminary tests (Table S5) did not show any degree of parthenogenesis. From the three *w* RNAi lines tested, one RNAi line did not show any degree of parthenogenesis, another showed a little parthenogenesis when the expression of the RNAi was driven but not in the control, and the last one showed a small degree of parthenogenesis in the control and when the expression of the RNAi was driven. Therefore, the small degree of parthenogenesis observed in two of the *w* RNAi stocks was considered to be a false positive result or the consequence of a background mutation. The *w* RNAi line that was positive when the expression was driven by Gal4 was added to the secondary screen.

iii) RNAi false positives

We found that the following RNAi lines used in the primary screen had a small degree of parthenogenesis without expression being driven by Gal4. These were the lines for *bam*, CG4496, CG10433, *CRMP*, *desat2*, *f*, and *FER*, and for the controls *Klp64D* and *w*. For all these experiments the RNAi line had fewer instances of parthenogenesis when crossed to the Gal4 drivers, and thus when the expression of the RNAi was induced. This indicates that the RNAi itself does not cause parthenogenesis but rather something in the background of the RNAi stocks. Since all these stocks were likely created from some of the same fly line(s) it is not unlikely that they all carry the same parthenogenesis inducing background mutation.

We tested the notion of a background mutation in the RNAi lines using the *bam* RNAi line. We crossed this line to 8 different *gal4* lines (*bam* (X), *bam* (III), *nos*, *MTB* (II), *MTB* (III), *actin* (*act*), and *ovarian tumour (otu)*) and found that some could drive a low level (0.1-0.2%) of parthenogenesis. However, this was less than or equal to the RNAi stock alone (0.2%). We also crossed the RNAi line to the *white* mutant that we found not to have any degree of parthenogenesis. Finally, we checked that the observed parthenogenesis could not be caused by the RNAi target gene by crossing the *bam* RNAi line to a mutant of *bam* and we found that this also reduced parthenogenesis. Therefore, we concluded that the only reasonable explanation is that the small degree of parthenogenesis is caused by a background mutation:

1) It is not the presence of the RNAi construct since many RNAi lines did not show any parthenogenesis.
2) It is not the expression of the RNAi since driving the expression did not enhance the degree of parthenogenesis.
3) It is not due to the *bam* mutation because crossing the RNAi line to a *bam* null mutant did not enhance the degree of parthenogenesis.

iv) Temperature selection for the screen

The first gene that we observed a reproducible amount of parthenogenesis with was the *bam* RNAi, therefore we carried out temperature optimisation with this RNAi line. This revealed the highest level of parthenogenesis at 18°C, moderate levels at 25°C, and no parthenogenesis at all at 29°C. We chose 25°C for our experiments as this would allow them to be completed in less than 3 months compared to 5 months at 18°C due to the difference in lifespan of the flies at these temperatures.

v) Statistical Analysis

All statistical analysis was done in R using Fisher's exact test. As control, we used the *w* mutant when comparing to other mutant alleles; the RNAi line in the absence of Gal4 driven expression, or the UAS-line without Gal4 driven expression. For ubiquitous expression or expression under an endogenous promotor (in the case of *GFP-polo*), we used Rcd4 expressed in a comparable manner as this showed no parthenogenesis. For the double gene variant screen, the indicated, listed controls were used; if there was no listed control then the single gene variant with the strongest phenotype was used as the control.

vi) Single gene variant screen

Summary:

We screened 28 candidates showing differential expression corresponding to a $\log_2$ fold change > 0.5 and padj < 0.05 from one or more of the three pairwise differential expression datasets (Table 2). We also screened 14 cell cycle/centrosome genes (Table S1) because they have long been implicated in causing parthenogenesis [18-20].

When the expression of 16 of these genes was either increased or decreased, we observed 0.1-0.4% parthenogenesis (Table 2, Table S1). This single gene variant screen required more extensive analysis since we did see a small amount of parthenogenesis in many of the controls listed above, suggesting that these were false positives resulting from a mutation present in the background of these fly lines.

Outcome – Data Table S7:

For clarity, we will consider the outcome for each gene listed in Data Table S7 to account for whether we deemed its mis-expression to contribute to parthenogenesis or not.

(1) *Asciz:* negative for parthenogenesis in the RNAi line tested

(2) *Bam:* negative for parthenogenesis, as outlined above.

(3) *c(2)M:* negative for parthenogenesis in the RNAi line tested.

*(4)* *Cad96Ca:* negative for parthenogenesis in the RNAi line tested.

(5) *CASK:* a deficiency, negative for parthenogenesis.

(6) *CG4329:* negative for parthenogenesis in the RNAi line tested.

(7) *CG4496*: 0.1% parthenogenesis only in the RNAi control and considered a false positive.

(8) *CG10433:* 0.1% parthenogenesis only in the RNAi control and considered a false positive.

(9) *CG17202:* a TE insertion allele, negative for parthenogenesis.

(10) *CG42808*: negative for parthenogenesis in the RNAi line tested.

(11) *chrb*: negative for parthenogenesis in the RNAi line tested.

(12) *CRMP*: 0.1% parthenogenesis only in the RNAi control and considered a false positive. However, the point mutant loss-of-function allele did show 0.2% parthenogenesis. This gene was therefore flagged for further testing.

(13) *desat1*: two RNAis were tested; one was negative for parthenogenesis and the other showed 0.2% parthenogenesis when RNAi was driven but not in the control. This was considered a positive result. An allele of *desat1* created by a transposable element (TE) insertion was also tested and showed the highest degree of parthenogenesis seen in our screen with 3 adult flies being produced. None of these flies were able to establish a parthenogenetic colony of flies. This gene was flagged for further testing.

(14) *desat2*: two RNAis were tested; one was negative for parthenogenesis and the other showed 0.1% parthenogenesis in the RNAi control and is therefore considered a false positive. An allele for *desat2* where the expression of the gene is reduced by a 16bp deletion 150bp upstream of the start site was also tested and had 0.3% parthenogenesis. This gene was flagged for further testing.

(15) *e(r)*: a TE insertion allele, negative for parthenogenesis.

(16) *eya*: overexpression was driven using two different UAS constructs - one showed a high degree of parthenogenesis (4 adult flies), however, this was only when the

expression construct was homozygous. When the gene was overexpressed then parthenogenesis was reduced. It was therefore deemed to be a false positive. The other construct was negative for parthenogenesis.

(17) *f*: 0.1% parthenogenesis in the RNAi control only and therefore deemed a false positive. However, the homozygous null mutant allele did show 0.1% parthenogenesis and so this gene was flagged for further testing.

(18) *FER*: 0.1% parthenogenesis only in the RNAi control and therefore considered a false positive. A TE insertion allele was negative for parthenogenesis.

(19) *gnu*: tested with one CRISPR allele and overexpression. The CRISPR allele showed 0.2% degree of parthenogenesis. Overexpression was negative for parthenogenesis. This gene was flagged for further testing.

(20) *ktub*: showed 0.1% parthenogenesis when RNAi was driven but not in the control. This was considered a positive result. A null allele of *ktub* was also tested and showed 0.4% degree of parthenogenesis. This gene was flagged for further testing.

(21) *msd1*: a TE insertion allele, negative for parthenogenesis.

(22) *Myc*: overexpression was driven using one UAS construct. It showed 0.2-0.3% parthenogenesis. This gene was flagged for further testing.

(23) *Nmnat*: overexpression showed 0.2% parthenogenesis when the overexpression construct was homozygous but there was no parthenogenesis when overexpressed, therefore it is likely a false positive.

(24) *pnt*: overexpression was negative for parthenogenesis.

(25) *Rcd4*: overexpression using two UAS constructs and two ubiquitous expression constructs was negative for parthenogenesis. These were also tested at two different temperatures (25°C, and 29°C).

(26) *Roc1a*: overexpression was negative for parthenogenesis; a TE insertion allele was also negative for parthenogenesis.

(27) *RpL10Ab*: a TE insertion allele, negative for parthenogenesis.

(28) *spir*: overexpression was negative for parthenogenesis.

The 14 Cell cycle/centrosome-related genes tested in the biased screen were:

(1) *ana2*: CRISPR allele was negative for parthenogenesis.

(2) *asl*: CRISPR allele showed 0.1% parthenogenesis. Overexpression was negative for parthenogenesis. This gene was flagged for further testing.

(3) *cnn*: A null allele showed 0.1% parthenogenesis. Since null CRISPR mutants in other pericentriolar material (PCM) genes (plp and asl) also gave similar results, we decided to screen those genes in the secondary screen.

(4) *cyclinE*: overexpression negative for parthenogenesis.

(5) *mr*: one CRISPR showed 0.1% parthenogenesis, and the other was negative. This gene was flagged for further testing.

(6) *Plk4/sak*: CRISPR null, a genome region duplication, and overexpression were all negative for parthenogenesis.

(7) *plp*: CRISPR allele showed 0.1% parthenogenesis. This gene was flagged for further testing.

(8) *plu*: two CRISPR alleles showed 0.1% parthenogenesis. Since Plu forms a complex with Png and Gnu [21], and *gnu* was also positive for parthenogenesis in the primary screen, we opted to only screen *gnu* in depth in the secondary screen and limited the screening of *plu*.

(9) *png*: two CRISPR alleles were negative for parthenogenesis.

(10) *polo*: a hypomorphic and a null allele were both negative for parthenogenesis. Overexpression showed 0.1% parthenogenesis. This gene was flagged for further testing.

(11) *rca1*: overexpression negative for parthenogenesis.

(12) *Sas-6*: CRISPR allele and UAS overexpression were negative for parthenogenesis. Ubiquitous overexpression showed 0.1% parthenogenesis but due to the inconsistent results and the fact that other centriole genes (*asl* and *Plk4*/*SAK*) did not cause parthenogenesis when overexpressed it was considered likely a false positive but was flagged for further testing.

(13) *slimb*: three CRISPR alleles showed 0.1-0.3% parthenogenesis. A deficiency and two loss of function alleles were negative. This gene was flagged for further testing.

(14) *tefu/ATM*: two RNAi lines and two loss of function alleles were negative for parthenogenesis.

vii) The double gene variant screen:

Summary:

We found combinations of 5 different variants that resulted in parthenogenesis (Table 3). These were *desat1* or *desat2* mutants, *slimb* mutant, and *polo* or *myc* overexpression constructs. These enhanced parthenogenesis from between 0.1-0.4% seen in the primary screen to 0.8-7.4%.

Detailed analysis:

We combined pairs of different alleles or expression constructs into the same fly stock and screened them for parthenogenesis. We specifically searched for combinations that enhanced parthenogenesis above the highest level of parthenogenesis observed in the single gene variant screen of 0.4% offspring produced. We tested combinations of the positive genes (Table 3, Data Table S8) in addition to *Plk4* and the putative false positives: *bam* RNAi, *eya* overexpression, *Klp64D* RNAi, *Trx-2* RNAi and mutant, and *w* RNAi. In addition, we also tested *Rcd4* overexpression as a negative control for double mutants. Here, we will go review the outcome for each group of genes and our considerations of whether it contributes to parthenogenesis or not.

(1) *asl* CRISPR: no enhancement of parthenogenesis when combined with polo overexpression and therefore considered to be a false positive.

(2) *bam* RNAi: resulted in parthenogenesis when homozygous (in the absence of Gal4) and in combination with overexpressed *GFP-polo.* Enhancement was not seen when RNAi was driven by Gal4. Thus, the effect upon parthenogenesis appears due to a background mutation in the *bam* RNAi line.

(3) *CRMP* mutant: when combined with 11 other mutants/RNAi lines/overexpression constructs, it showed no enhancement of parthenogenesis but a slight decrease. We therefore determined CRMP to not enhance parthenogenesis and likely to be a false positive.

(4) *desat1* RNAi: when combined with 11 other mutants/RNAis/overexpression, there was only one combination that showed any enhancement of parthenogenesis and that was with the *Klp64D* RNAi, which is not caused by the RNAi itself but by a background mutation. Thus, *desat1* RNAi does not appear effective at inducing parthenogenesis with the caveats indicated above for RNAi knock-down in the germline.

(5) *desat1* mutant: when combined with 13 other mutants/RNAi lines/overexpression constructs, two showed significant enhancement – heterozygous *desat1* with *polo* overexpression enhanced parthenogenesis from 0.4% to 5.3%, a greater than 10-fold increase. A combination of *desat1* with the *Trx-2* also enhanced parthenogenesis to

1.3%. However, we considered this combination likely to be a false positive (discussed below).

(6) *desat2* RNAi: when combined with 12 other mutants/RNAi lines/overexpression constructs, the combination with *Trx-2* RNAi showed enhancement of parthenogenesis and this appears not to be not caused by the targeting of *Trx-2* but by a possible off-target effect. We deemed that the *desat2* RNAi was not effective at enhancing parthenogenesis.

(7) *desat2* mutant: when combined with 14 other mutants/RNAi lines/overexpression constructs, the combination of heterozygous *desat2* with *GFP-polo* overexpression enhanced parthenogenesis from 0.4% to 7.4%, over 10-fold; and the combination of *desat2* with *slimb* enhanced parthenogenesis to 1.2%.

(8) *eya* overexpression: when combined with 12 other mutants/RNAi lines/overexpression constructs, enhancement of parthenogenesis to 0.7% was only observed in combination with *ktub*. No other combination showed enhancement of parthenogenesis therefore we concluded that *eya* was likely a false positive.

(9) *f* mutant: when combined with 11 other mutants/RNAi lines/overexpression constructs there was an approximate increase in parthenogenesis (0.1% to 0.9%) in combination with *polo* overexpression. We deem this to be a modest enhancer of parthenogenesis.

(10) *gnu* CRISPR: when combined with 5 other mutants there was no enhancement of parthenogenesis, and it was therefore deemed to be a false positive.

(11) *Klp64D* RNAi: when combined with 11 other mutants/RNAi lines/overexpression constructs, we saw enhancement of parthenogenesis with *desat1* RNAi, which is not caused by the RNAi itself but by a background mutation. Thus, *Klp64D* RNAi appears not to be effective at inducing parthenogenesis.

(12) *ktub* mutant: when combined with 13 other mutants/RNAi lines/overexpression constructs, there was a modest increase in parthenogenesis (from 0.4% to 0.6-0.8%) with *eya* overexpression, *polo* overexpression, and with the *slimb* mutant. This increase does not appear significant. When combined with *desat1* there was no enhancement, and the flies were not fertile when combined with *desat2*. We therefore deem *ktub* not to enhance parthenogenesis or to be a false positive.

(13) *mr* CRISPR allele: when combined with *polo* overexpression there was no enhancement of parthenogenesis suggesting it to be false positive.

(14) *Myc* overexpression: in combination with *GFP-polo* overexpression there was 0.6% parthenogenesis but a decrease in combination with *desat1* and *desat2*. As there was zero parthenogenesis in the *UAS-Myc* stock when not combined with a gal4 stock, we concluded that this is a positive result. It is likely that *Myc* overexpression is too deleterious when combined with other mutants because we are only able to drive *Myc* expression very strongly with the UAS/gal4 system and therefore could not mimic the slight overexpression seen in *D. mercatorum*. Alternatively, changes in *desat1* and *desat2* expression could lie downstream of Myc and this may mask any enhancement.

(15) *plp* CRISPR allele: we found no enhancement when combined with *polo* overexpression or the *desat1* mutant and therefore deem *plp* a false positive.

(16) *plu* CRISPR allele: we found no enhancement when combined with the *desat2* mutant and therefore deem *plu* a false positive.

(17) *Plk4/Sak* CRISPR allele: *plk4* mutant did not show any degree of parthenogenesis alone but when combined with *slimb*, it enhanced parthenogenesis to 0.8%. The interpretation of this result is uncertain. Plk4/SAK is targeted by *slimb* for destruction and in the absence of Slimb, Plk4 levels are elevated leading to

excessive centriole biogenesis [22, 23]. It might be anticipated that inactivating one copy of *slimb* and removing one copy of *Plk4* would be neutral for centriole biogenesis suggesting some other functional route. This finding therefore needs further investigation.

(18) *GFP-Polo* overexpression: alone *GFP-polo* overexpression results in only 0.1% parthenogenesis but when combined with 20 other mutants/RNAi lines/overexpression constructs, there was an enhancement of parthenogenesis observed for nearly all combinations. Enhancement is particularly notable with *bam* RNAi, *desat1* mutant, *desat2* mutant, *ktub* mutant, and *slimb* mutant. The combination with *bam* RNAi and *Trx-2* RNAi is likely to be a false positives because the increase to 1.6-2.6% was only seen when the RNAi construct was homozygous and not when it was expressed with a Gal4. The *desat1/2* and *ktub* mutant are discussed above. The *slimb* mutant enhanced parthenogenesis to 3.1%. Taken together *polo* overexpression was the strongest enhancer of parthenogenesis observed in our screen.

(19) *Rcd4* overexpression: with *bam* RNAi there was no enhancement of parthenogenesis, and it was therefore considered an appropriate negative control for the double mutant screen.

(20) *Sas-6* overexpression: we found no enhancement when combined with the *desat1* mutant and therefore deem *Sas-6* a false positive.

(21) *slimb* mutants: when in 12 mutant combinations, the level of parthenogenesis was enhanced from 0.3% to 0.6-3.1% by only *desat2*, *ktub*, and *plk4* mutants, and by *polo* overexpression. All other combinations are discussed above.

(22) *Trx-2* RNAi: enhancement of parthenogenesis was seen in combination with *desat2* RNAi and *polo* overexpression. In combination with *polo* overexpression,

parthenogenesis decreased when the RNAi construct was expressed using Gal4 driver or when heterozygous, therefore the increase in parthenogenesis is not likely cause by the RNAi or the reduced expression of *Trx-2* because we only observed a decrease in phenotype when combined with the *Trx-2* mutant. When the *Trx-2* RNAi was combined with *desat2* RNAi we observed 0.2% parthenogenesis, which increased to 2.4% when the RNAis were expressed with the gal4. We believe that since there are no other combinations of *desat2* and *Trx-2* gave a similar phenotype that this was a combination of favourable background mutations that could be enhanced by *desat2* knockdown.

(23) *Trx-2* mutant: this showed enhancement of parthenogenesis (from zero to 1.3%) when homozygous and in combination with heterozygous *desat1* mutant. Considering that we only saw enhanced levels of parthenogenesis in this specific combination compared to the 10 other combinations that either showed no parthenogenesis or no enhancement, we consider this to only be a specific enhancer for heterozygous *desat1* or a false positive.

(24) *w* RNAi: when combined with 11 other mutants we saw no enhancement of parthenogenesis suggesting it to be a false positive.

The conclusion from this screen was that the *desat1* or *desat2* mutants, *slimb* mutant, and *polo* or *Myc* overexpression led to parthenogenesis.

The genetic changes that underly parthenogenesis

We did not find any striking differences at the *desat* locus between the parthenogenetic and sexual genomes. In the first *desat1* exon there are two C to T transversions, one C to G transversion in the second exon, and a C to T transversion in the fourth exon. None of these mutations change the protein sequence, which is identical in the sexually reproducing and parthenogenetic strains. In the first, second, and third introns there are many transitions, transversions, and small deletions. None of these seem likely to be of consequence to gene expression. We detected two deletions upstream of the gene with potential to lead to its differential expression, a 4bp deletion 60bp upstream and a 20bp deletion 712bp upstream of the *desat1* ORF (Fig. S7A). In *D. melanogaster* the untranslated region (UTR) extends up to 5kbp upstream of the start site, therefore it is possible that these mutations could affect transcription of *desat1*.

Although *desat2* showed very high differential expression, and we found no substantial changes to the *desat2* locus there, the coding sequence remained exactly the same between the sexual and parthenogenetic genomes. There is a 4 bp deletion in the first intron which is unlikely to interfere with expression (Fig. S7B). It is more likely that there is a change to an unknown upstream regulator of transcription resulting in the change in expression for *desat2*. Based on our finding that there were limited genomic changes near this *desat* locus, that both genes had a similar phenotype, they do not enhance the phenotype of each other, and that only *desat2* was significantly differentially expressed, we propose that reduced overlapping functions of these two genes contributes to parthenogenesis.

Increasing *polo* levels also caused parthenogenesis although this gene was not differentially expressed in the germline. There was a 113bp deletion 1130bp upstream of the start site for the *polo* gene that was unlikely to affect gene expression and the coding region was a perfect match between the two genomes (Fig. 7C). This accords with transcriptomics finding that there is no differential expression and supports our notion assertion that Polo could be a downstream effector of primary mutations or a result of *Myc* overexpression.

There were minor changes to the genome regions for *slimb* (Fig. 7D) In the *slimb* sixth exon there is a T to C transition which does not influence the coding sequence, and hence the protein is identical. In the introns there are many transitions, transversions, and small deletions. Lack of differential expression suggests that none of these are likely to be of consequence to the expression this gene.

**Supplementary Figures:**

**Fig. S1**. **Gene distribution and sequencing read coverage for the *D. mercatorum* genomes.** (**A**) Genes per contig for the sexually reproducing *D. mercatorum* genome. (B) Genes per contig for the parthenogenetic *D. mercatorum* genome. (C) Plot of the read coverage across the sexually reproducing *D. mercatorum* genome. (D) Plot of the read coverage across the parthenogenetic *D. mercatorum* genome.

**Fig. S2**. **Genome alignment of parthenogenetic *D. mercatorum* against *D. melanogaster.*** Parthenogenetic *D. mercatorum* genome assembly aligned against the *D. melanogaster* reference genome (release 6). The purple dots or lines represent sequences matching against the forward strand and the blue against the reverse**.**

**Fig. S3**. **Match of *D. mercatorum* contigs to *D. melanogaster* chromosome arms.** (**A**) Chromosome arm assignment for the sexual *D. mercatorum* genome based on k-mer mapping using nucmer. (**B**) Chromosome arm assignment for the parthenogenetic *D. mercatorum* genome based on k-mer mapping using nucmer.

**Fig. S4**. **Transcriptomics analysis.** (**A**) Expressed genes by contig for the sexual *D. mercatorum* genome showing that gene expression is distributed across the contigs. (**B**) Expressed genes by contig for the parthenogenetic *D. mercatorum* genome showing that the gene expression is distributed across the contigs. (**C**) Upset plot of the number of genes that were differentially expressed based on comparison set - from left to right: the genes overlapping between all three pairwise comparisons, the combinations of the two different pairwise comparison (red bars), and the number of genes in single set comparisons (black bars). Candidate genes at Padj<0.05, significant according to Super Exact Test. (**D**) Gene ontology (GO) analysis for all pairwise transcriptomics comparisons. GO enrichment (Fisher's exact test) and gene set enrichment analysis (GSEA, Kolmogorov-Smirnov test) using the *D. melanogaster* homologues. The frequency depicts how often a GO category was below this FDR (false discovery rate) threshold. (**E**) Volcano plots $\log_2$ fold change vs $\log_{10}$ *p*adj for the parthenogenetic vs sexual transcriptomes, partially parthenogenetic vs parthenogenetic transcriptomes, and partially parthenogenetic vs sexual transcriptomes. The screened genes are indicated in the legend.

**Fig. S5**. **Differential expression plots for all genes that were screened.** (**A**) Differential expression between all three transcriptomics library comparisons for the candidate genes that were selected from the differential expression analysis. (**B**) Differential expression between all three transcriptomics library comparisons for the candidate cell cycle/centrosome genes that were screened, excepting tefu/ATM whose expression could not be detected in *D. mercatorum* eggs. (**C**) Differential expression between all control genes that were screened, except *white* whose expression could not be detected in *D. mercatorum* eggs.

**Fig. S6**. **CRISPR alleles created for the parthenogenesis screen.** (**A**) *ana2* CRISPR design and resulting allele. (**B**) *asl* CRISPR design and resulting allele. (**C**) *gnu* CRISPR design and resulting allele. (**D**) *mr* CRISPR design and resulting alleles. (**E**) *plk4* CRISPR design and resulting allele. (**F**) *Plp* CRISPR design and resulting allele. (**G**) *plu* CRISPR design and resulting alleles. (**H**) *png* CRISPR design and resulting alleles. (**I**) *Sas-6* CRISPR design and resulting allele. (**J**) *slimb* CRISPR design and resulting alleles. On the gene representation the grey boxes are the untranslated region (UTR) and the blue boxes are the exons. The gRNA sequence is given in blue and the protospacer adjacent motif in pink. The cut site is indicated with a pink arrow. The start and the end site are in bold, and any added bases are black and bolded.

**Fig. S7**. **Genomic loci of the genes that caused parthenogenesis.** (**A**) Genomic changes at the *desat1* locus. (**B**) Genomic changes at the *desat2* locus. (**C**) Genomic changes at the *polo* locus. (**D**) Genomic changes at the *slimb* locus. (**E**) Genomic and coding changes at the *Myc* locus.

**Fig. S8**. **Myc protein sequence comparison between species.** (**A**) Comparison between mouse, human, mosquito, *D. melanogaster,* sexual *D. mercatorum*, and parthenogenetic *D. mercatorum*, and aligned with CLUSTAL O(1.2.4) multiple sequence alignment tool.

**Fig. S9**. **Parthenogenetic and sexual *D. mercatorum* embryos.** (**A**) Fertilized sexually reproduced embryo that has entered the mitotic nuclear divisions. (**B**) Partially parthenogenetic embryo that has completed meiosis, showing the three polar bodies beginning to form an aggregate. (**C**) Partially parthenogenetic embryo that has initiated mitosis. (**D**) Partially parthenogenetic embryo that has entered the mitotic cell divisions; there are 11 prophase nuclei but not all are visible on this image. (**E**) Parthenogenetic embryo that has completed meiosis, showing the 4 products of meiosis. (**F**) Parthenogenetic embryo that has initiated mitosis. (**G**) Parthenogenetic embryo that has entered the mitotic cell divisions. The nuclei are marked with asterisks and the sperm tail is marked with a lined arrow. The scale is 10μm.

**Fig. S10**. **Wild-type *D. melanogaster* eggs and embryos**. (**A**) Unfertilized embryo that has completed meiosis, showing the polar body aggregate. (**B**) Unfertilized embryo that has initiated mitosis. (**C**) Fertilized embryo that has not yet initiated mitosis. Note the sperm tail marked with acetylated tubulin staining. (**D**) Fertilized embryo that is undergoing the second round of mitosis (in anaphase). Nuclei are marked with asterisks. The polar body aggregate when shown with other nuclei is marked with an arrowhead. The sperm tail is marked with a lined arrow. The scale bar is 10μm.

**Fig. S11**. **Parthenogenetic and sexually reproducing *D. melanogaster* embryos**. (**A**) *GFP-polo* embryo that has completed meiosis, showing the typical polar body aggregate. (**B**) *GFP-polo* embryo that has initiated mitosis. (**C**) *desat2* embryo that has completed meiosis, showing an abnormal polar body aggregate. (**D**) *desat2* embryo that has initiated mitosis (**E**) *GFP-polo*$^{4+}$*; desat2*$^{-/+}$ embryo that has completed meiosis, having the abnormal polar body characteristic of *desat1/2* mutants. (**F**) *GFP-polo*$^{4+}$*; desat2*$^{-/+}$ embryo that has initiated mitosis showing the centrosome-like GFP-Polo puncta. (**G**) *GFP-polo*$^{4+}$*; desat2*$^{-/+}$ embryo that has entered the mitotic nuclear divisions, having 4 nuclei in metaphase. Nuclei are marked with asterisks. The scale bar is 10μm.

**Supplementary Tables:**

**Table S1**. **Biased screen of cell cycle and centrosome genes.** Gene function was attributed from flybase.org and the screen was performed with the indicated genetic tools. The percent refers to the number of parthenogenetic offspring produced relative to the number of virgin females screened. The *p* value for the proportion of parthenogenetic offspring versus virgin females screened was calculated using the Fisher's exact test.

**Table S2**. **Single gene variant screen controls**. Negative controls that were not differentially expressed. Screens were performed with the indicated genetic tools. The percent refers to the number of parthenogenetic offspring produced relative to the number of virgin females screened.

**Table S3**. **Parthenogenetically produced offspring backcrossed to males**. The parthenogenetically produced offspring from the genetically induced *D. melanogaster* and naturally produced *D. mercatorum* from the partially parthenogenetic stock were back crossed to the parental line to determine if they were able to produce male and female in their F2 generation. The backcross was marked as successful or not.

**Supplementary Data Tables:**

**Data Table S1**: **Parthenogenesis in *D. mercatorum***. Baseline experiments that were carried out to determine expectation when screening for parthenogenesis in *D. mercatorum* and to aid in the selection of which strains to study in more detail.

**Data Table S2**: **Hybridisation experiments between different *D. mercatorum* strains**. Between 3-10 females and males from the strains listed below were crossed in equal numbers and maintained together on fresh food for the duration of their life. If offspring were produced, we checked for the presence of at least 3 male offspring, as an indication that mating had occurred, and the cross was scored 'yes'. If no male offspring were produced, these experiments were scored 'no'. These experiments were repeated 3 or 4 times. If F1 were generated, then they were flipped into a new tube and checked for their ability to produce offspring.

**Data Table S3**: **Summary tables of differential gene expression (DE) analysis**. The three different transcriptomic comparisons are indicated together with overlapping genes that are differentially expressed in all datasets.

**Data Table S4**: **Parthenogenesis in different *Drosophila* species**. These experiments were carried out to gain understanding of expectations in screens for variable levels of parthenogenesis and estimate the optimal number of virgins to collect for each experiment.

**Data Table S5**: **Parthenogenesis in different *D. melanogaster* strains**. These experiments were carried out to confirm that *D. melanogaster* is not naturally parthenogenetic.

**Data Table S6**: **CRISPR mutant information**. CRISPR gRNA sequences, sequencing primers, resulting CRISPR allele genotype, viability, rescue data, and complementation.

**Data Table S7**: **Single gene variant screen**. Tracking of the temperature at which flies were maintained; the number of crosses, if applicable; the number of flies collected; the maximum average lifespan per batch; the average maternal age of parthenogenesis; proportion of life completed at the age of parthenogenesis; the maximum developmental stage reached by parthenogenetic offspring; proportion and percent of offspring produced compared to the number of flies screened; and any additional observations about any adult flies generated; *p* value when appropriate; and the source of the fly stock.

**Data Table S8**: **Double gene variant screen**. Tracking of the temperature at which flies were maintained; the number of crosses, if applicable; the number of flies collected; the maximum average lifespan per batch; the average maternal age of parthenogenesis; proportion of life completed at the age of parthenogenesis; the maximum developmental stage reached by parthenogenetic offspring; proportion and percent of offspring produced compared to the overall number of flies screened; any additional observations about any adult flies generated; and *p* value when appropriate.
