## SupplementaryFigures+Tables for "Virgin Birth: A genetic basis for facultative parthenogenesis"

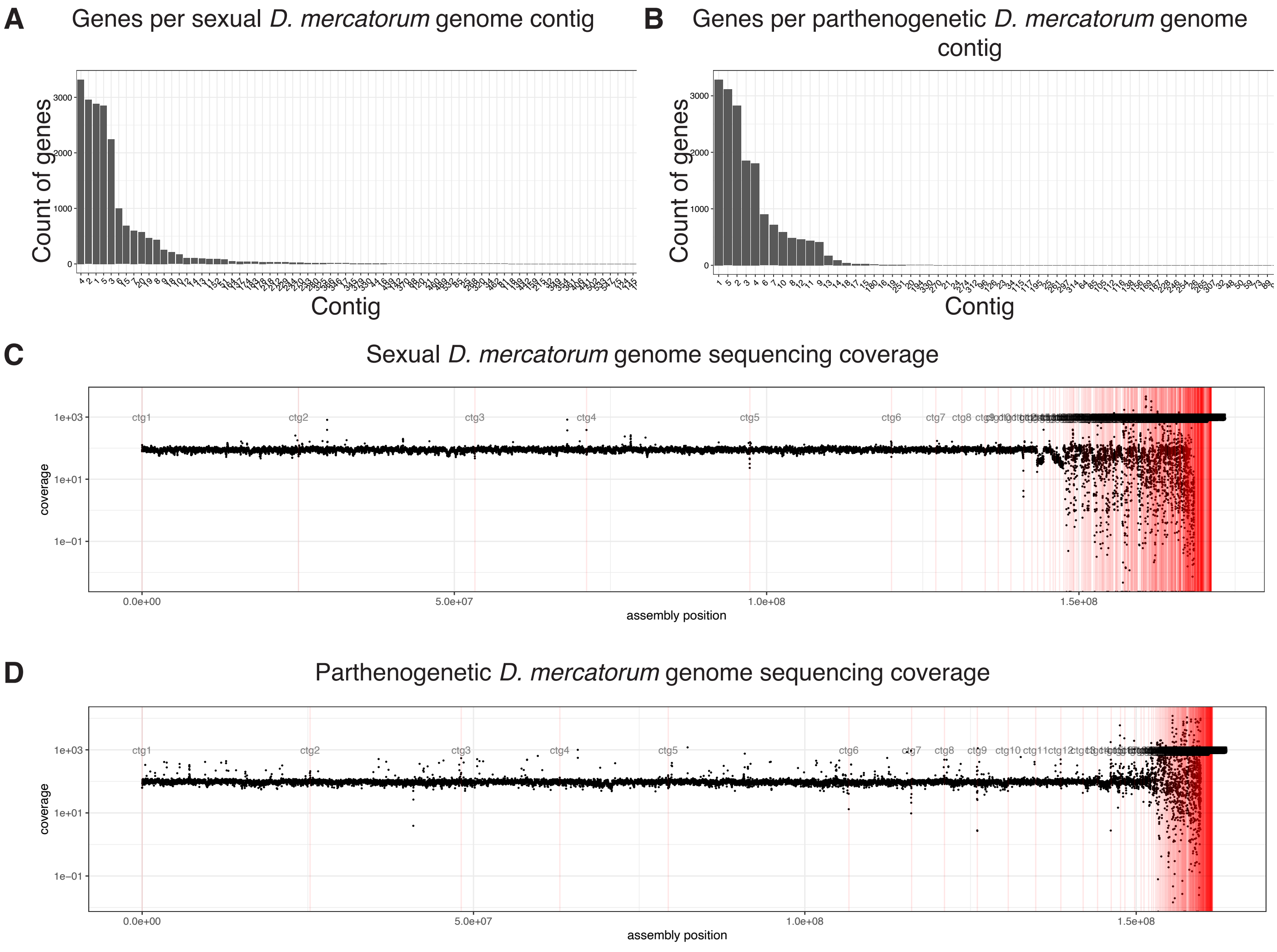
A
Genes per sexual D. mercatorum genome contig
Count of genes
3000
2000
1000
0
Contig
B
Genes per parthenogenetic D. mercatorum genome contig
Count of genes
3000
2000
1000
0
Contig
C
Sexual D. mercatorum genome sequencing coverage
coverage
1e+03
1e+01
1e−01
ctg1
ctg2
ctg3
ctg4
ctg5
ctg6
ctg7
ctg8
0.0e+00
5.0e+07
1.0e+08
1.5e+08
assembly position
D
Parthenogenetic D. mercatorum genome sequencing coverage
coverage
1e+03
1e+01
1e−01
ctg1
ctg2
ctg3
ctg4
ctg5
ctg6
ctg7
ctg8
ctg9
ctg10
ctg11
ctg12
0.0e+00
5.0e+07
1.0e+08
1.5e+08
assembly position

**Figure S1:**

**A**

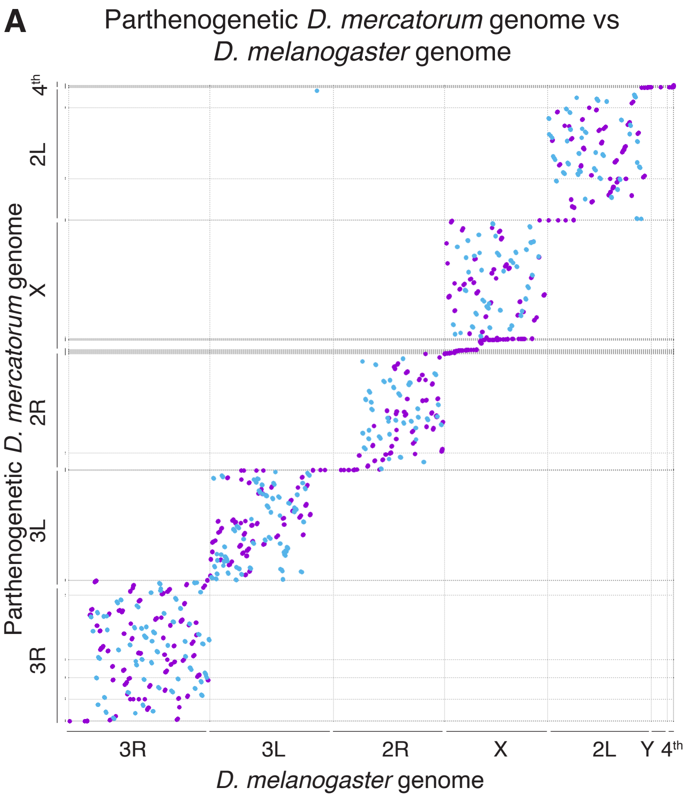

Parthenogenetic *D. mercatorum* genome vs
*D. melanogaster* genome
Parthenogenetic *D. mercatorum* genome
4th
2L
X
2R
3L
3R
3R
3L
2R
X
2L
Y
4th
*D. melanogaster* genome


**Figure S2:**

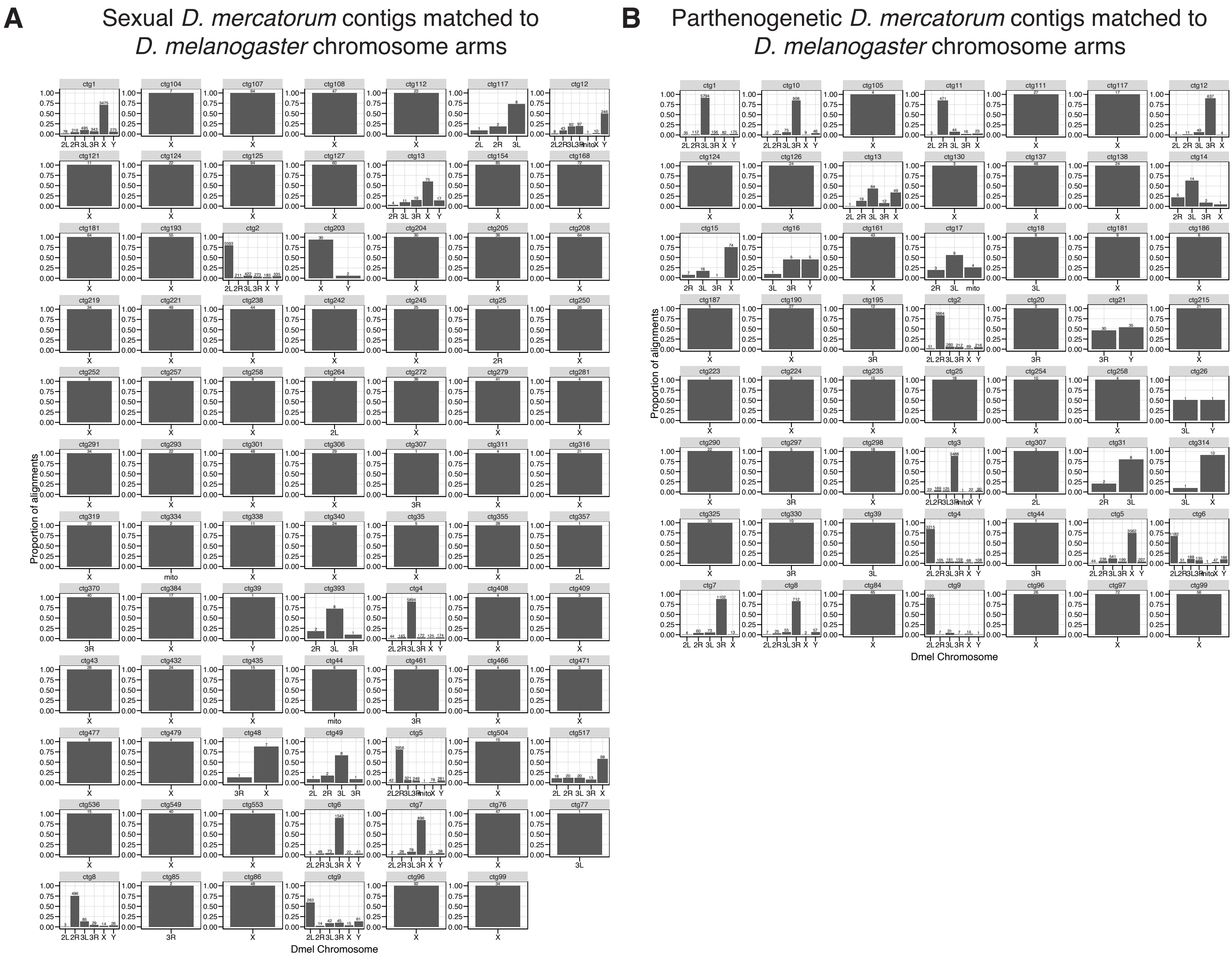
A
Sexual D. mercatorum contigs matched to D. melanogaster chromosome arms
B
Parthenogenetic D. mercatorum contigs matched to D. melanogaster chromosome arms
Proportion of alignments
Dmel Chromosome

**Figure S3:**

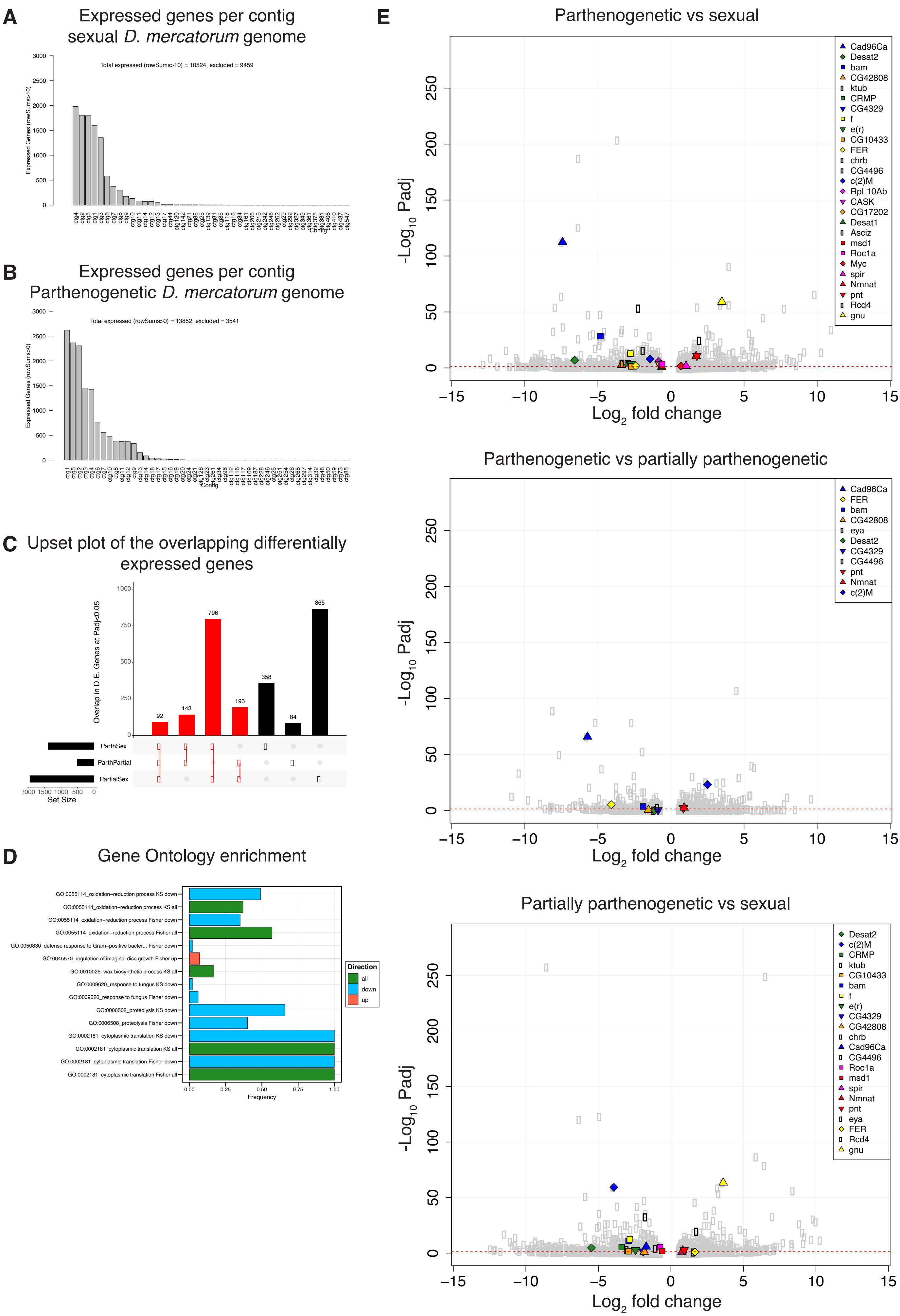
A
Expressed genes per contig
sexual D. mercatorum genome
Total expressed (rowSums>10) = 10524, excluded = 9459
Expressed Genes (rowSums>10)
Contig
B
Expressed genes per contig
Parthenogenetic D. mercatorum genome
Total expressed (rowSums>0) = 13852, excluded = 3541
Expressed Genes (rowSums>0)
Contig
C
Upset plot of the overlapping differentially expressed genes
Overlap in D.E. Genes at Padj<0.05
92
143
796
193
358
84
865
ParthSex
ParthPartial
PartialSex
Set Size
D
Gene Ontology enrichment
GO:0055114_oxidation−reduction process KS down
GO:0055114_oxidation−reduction process KS all
GO:0055114_oxidation−reduction process Fisher down
GO:0055114_oxidation−reduction process Fisher all
GO:0050830_defense response to Gram−positive bacter... Fisher down
GO:0045570_regulation of imaginal disc growth Fisher up
GO:0010025_wax biosynthetic process KS all
GO:0009620_response to fungus KS down
GO:0009620_response to fungus Fisher down
GO:0006508_proteolysis KS down
GO:0006508_proteolysis Fisher down
GO:0002181_cytoplasmic translation KS down
GO:0002181_cytoplasmic translation KS all
GO:0002181_cytoplasmic translation Fisher down
GO:0002181_cytoplasmic translation Fisher all
Frequency
Direction
all
down
up
E
Parthenogenetic vs sexual
-Log10 Padj
Log2 fold change
Cad96Ca
Desat2
bam
CG42808
ktub
CRMP
CG4329
f
e(r)
CG10433
FER
chrb
CG4496
c(2)M
RpL10Ab
CASK
CG17202
Desat1
Asciz
msd1
Roc1a
Myc
spir
Nmnat
pnt
Rcd4
gnu
Parthenogenetic vs partially parthenogenetic
-Log10 Padj
Log2 fold change
Cad96Ca
FER
bam
CG42808
eya
Desat2
CG4329
CG4496
pnt
Nmnat
c(2)M
Partially parthenogenetic vs sexual
-Log10 Padj
Log2 fold change
Desat2
c(2)M
CRMP
ktub
CG10433
bam
f
e(r)
CG4329
CG42808
chrb
Cad96Ca
CG4496
Roc1a
msd1
spir
Nmnat
pnt
eya
FER
Rcd4
gnu

**Figure S4:**

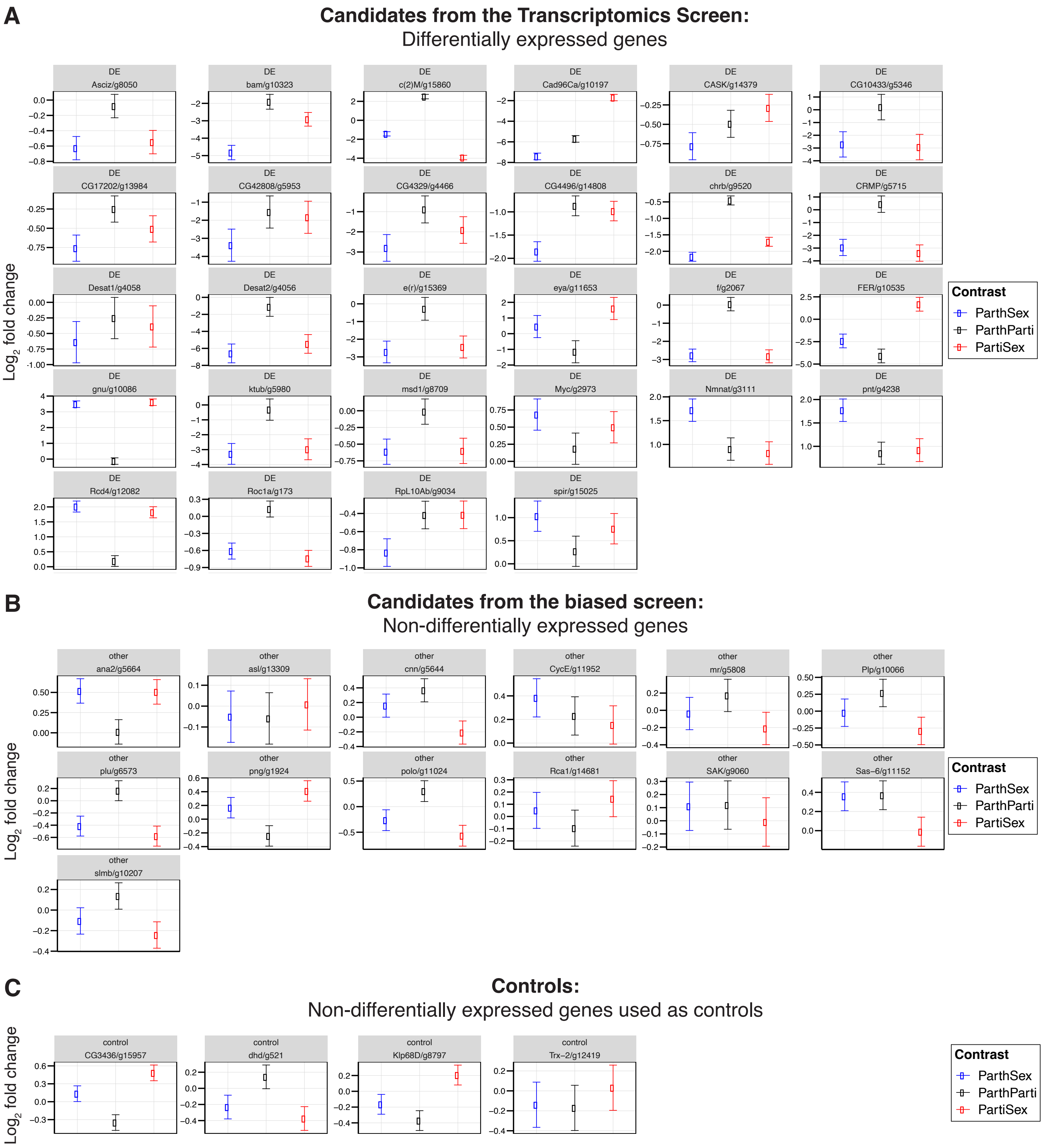
A
Candidates from the Transcriptomics Screen:
Differentially expressed genes
Log2 fold change
DE Asciz/g8050
DE bam/g10323
DE c(2)M/g15860
DE Cad96Ca/g10197
DE CASK/g14379
DE CG10433/g5346
DE CG17202/g13984
DE CG42808/g5953
DE CG4329/g4466
DE CG4496/g14808
DE chrb/g9520
DE CRMP/g5715
DE Desat1/g4058
DE Desat2/g4056
DE e(r)/g15369
DE eya/g11653
DE f/g2067
DE FER/g10535
DE gnu/g10086
DE ktub/g5980
DE msd1/g8709
DE Myc/g2973
DE Nmnat/g3111
DE pnt/g4238
DE Rcd4/g12082
DE Roc1a/g173
DE RpL10Ab/g9034
DE spir/g15025
Contrast
ParthSex
ParthParti
PartiSex
B
Candidates from the biased screen:
Non-differentially expressed genes
Log2 fold change
other ana2/g5664
other asl/g13309
other cnn/g5644
other CycE/g11952
other mr/g5808
other Plp/g10066
other plu/g6573
other png/g1924
other polo/g11024
other Rca1/g14681
other SAK/g9060
other Sas-6/g11152
other slmb/g10207
Contrast
ParthSex
ParthParti
PartiSex
C
Controls:
Non-differentially expressed genes used as controls
Log2 fold change
control CG3436/g15957
control dhd/g521
control Klp68D/g8797
control Trx-2/g12419
Contrast
ParthSex
ParthParti
PartiSex

**Figure S5:**

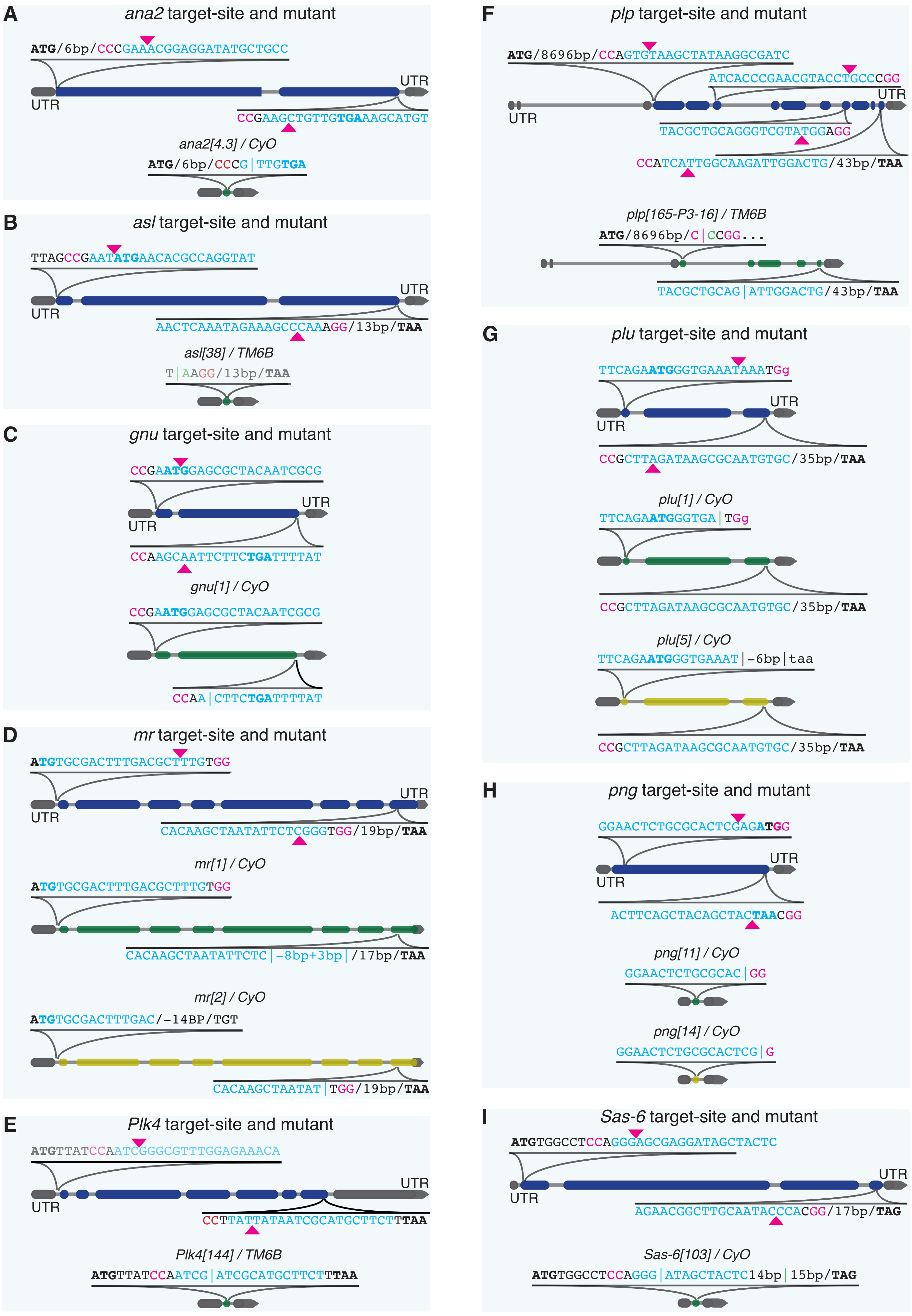
A
ana2 target-site and mutant
ATG/6bp/CCCGAAACGGAGGATATGCTGCC
UTR
UTR
CCGAAGCTGTTGTGAAAGCATGT
ana2[4.3] / CyO
ATG/6bp/CCCG|TTGTGA
B
asl target-site and mutant
TTAGCCGAATATGAACACGCCAGGTAT
UTR
UTR
AACTCAAATAGAAAGCCCAAAGG/13bp/TAA
asl[38] / TM6B
T|AAGG/13bp/TAA
C
gnu target-site and mutant
CCGAATGGAGCGCTACAATCGCG
UTR
UTR
CCAAGCAATTCTTCTGATTTTAT
gnu[1] / CyO
CCGAATGGAGCGCTACAATCGCG
CCAA|CTTCTGATTTTAT
D
mr target-site and mutant
ATGTGCGACTTTGACGCTTTGTGG
UTR
UTR
CACAAGCTAATATTCTCGGGTGG/19bp/TAA
mr[1] / CyO
ATGTGCGACTTTGACGCTTTGTGG
CACAAGCTAATATTCTC|-8bp+3bp|/17bp/TAA
mr[2] / CyO
ATGTGCGACTTTGAC/-14BP/TGT
CACAAGCTAATAT|TGG/19bp/TAA
E
Plk4 target-site and mutant
ATGTTATCCAATCGGGCGTTTGGAGAAACA
UTR
UTR
CCTTATTATAATCGCATGCTTCTTTAA
Plk4[144] / TM6B
ATGTTATCCAATCG|ATCGCATGCTTCTTTAA
F
plp target-site and mutant
ATG/8696bp/CCAGTGTAAGCTATAAGGCGATC
ATCACCCGAACGTACCTGCCCGG
UTR
UTR
TACGCTGCAGGGTCGTATGGAGG
CCATCATTGGCAAGATTGGACTG/43bp/TAA
plp[165-P3-16] / TM6B
ATG/8696bp/C|CCGG...
TACGCTGCAG|ATTGGACTG/43bp/TAA
G
plu target-site and mutant
TTCAGAATGGGTGAAATAAATGg
UTR
UTR
CCGCTTAGATAAGCGCAATGTGC/35bp/TAA
plu[1] / CyO
TTCAGAATGGGTGA|TGg
CCGCTTAGATAAGCGCAATGTGC/35bp/TAA
plu[5] / CyO
TTCAGAATGGGTGAAAT|-6bp|taa
CCGCTTAGATAAGCGCAATGTGC/35bp/TAA
H
png target-site and mutant
GGAACTCTGCGCACTCGAGATGG
UTR
UTR
ACTTCAGCTACAGCTACTAACGG
png[11] / CyO
GGAACTCTGCGCAC|GG
png[14] / CyO
GGAACTCTGCGCACTCG|G
I
Sas-6 target-site and mutant
ATGTGGCCTCCAGGGAGCGAGGATAGCTACTC
UTR
UTR
AGAACGGCTTGCAATACCCACGG/17bp/TAG
Sas-6[103] / CyO
ATGTGGCCTCCAGGG|ATAGCTACTC14bp|15bp/TAG

J

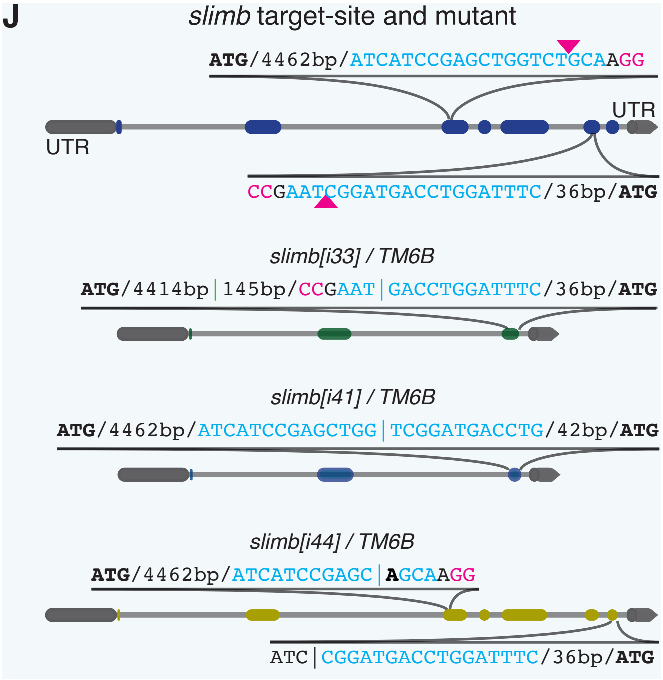

*slimb* target-site and mutant
ATG/4462bp/ATCATCCGAGCTGGTCTGCAAGG
UTR
UTR
CCGAATCGGATGACCTGGATTTC/36bp/ATG
*slimb[i33] / TM6B*
ATG/4414bp|145bp/CCGAAT|GACCTGGATTTC/36bp/ATG
*slimb[i41] / TM6B*
ATG/4462bp/ATCATCCGAGCTGG|TCGGATGACCTG/42bp/ATG
*slimb[i44] / TM6B*
ATG/4462bp/ATCATCCGAGC|AGCAAGG
ATC|CGGATGACCTGGATTTC/36bp/ATG


**Figure S6:**

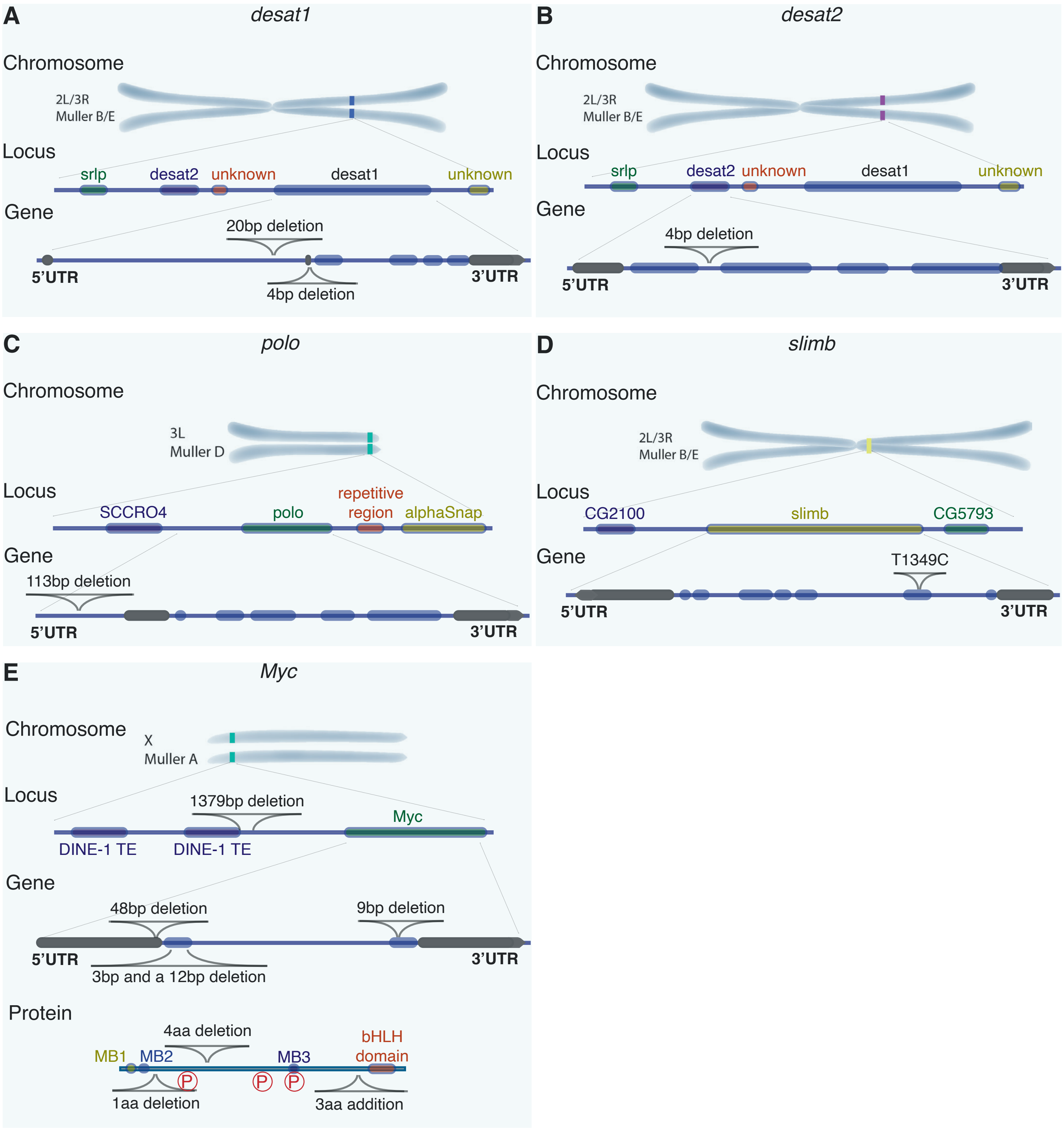
A
desat1
Chromosome
2L/3R
Muller B/E
Locus
srlp
desat2
unknown
desat1
unknown
Gene
20bp deletion
5'UTR
3'UTR
4bp deletion
B
desat2
Chromosome
2L/3R
Muller B/E
Locus
srlp
desat2
unknown
desat1
unknown
Gene
4bp deletion
5'UTR
3'UTR
C
polo
Chromosome
3L
Muller D
Locus
repetitive region
SCCRO4
polo
alphaSnap
Gene
113bp deletion
5'UTR
3'UTR
D
slimb
Chromosome
2L/3R
Muller B/E
Locus
CG2100
slimb
CG5793
Gene
T1349C
5'UTR
3'UTR
E
Myc
Chromosome
X
Muller A
Locus
1379bp deletion
Myc
DINE-1 TE
DINE-1 TE
Gene
48bp deletion
9bp deletion
5'UTR
3'UTR
3bp and a 12bp deletion
Protein
4aa deletion
bHLH domain
MB1
MB2
MB3
P
P
P
1aa deletion
3aa addition

**Figure S7:**

**A** Myc protein sequence comparison between mouse, human, mosquito, *D. melanogaster*, sexual *D. mercatorum*, and parthenogenetic *D. mercatorum*

```
mouse                           ------------------------------------------------------------   0
human                           ------------------------------------------------------------   0
mosquito                        MVSIKQEPSCWDDIKTISIKQELSNWDDSH--NMDIDWEQDIGIQFMDLPTSEFLTSAVE   58
Dmelanogaster                   -----------MAL---YRSDPYSIMDDQLFSNISIFDMDNDLYDMDKLLSSSTIQSDLE   46
Sexual_Dmercatorum              -----MTTACSSGI---CISGEFDLMDEMGFDLLE-FNVQDIGY------RLPSIQNDLE   45
parthenogenetic_Dmercatorum     -----MTTACSSGI---CISGEFDLMDEMGFDLLE-FNVQDIGY------RLPSIQNDLE   45

                                     Myc box 1              Myc box 2
mouse                           ------------------------------------------------------------   0
human                           ------------------------------------------------------------   0
mosquito                        LEQTYGSATCPANGWEQPASSKTQIRNHDCMWSGTCFDQSHPGKMGCGTNHGPANTTQDQ   118
Dmelanogaster                   KIEDMESV-F--QDYDLEEDMKPEIRNIDCMWPAMSS--CLTSGNGNGIE----------   91
Sexual_Dmercatorum              KIAAEHAHNMNSLALADDFDIKPEIRNGDCMWSAFGS--SANGGVNGANNNNNNNNSSNI   103
parthenogenetic_Dmercatorum     KIAAEHAHNMNSLALADDFDIKPEIRNGDCMWSAFGS--SANGGVNGANNNN-NNNSSNI   102
                                                                                1 aa deletion

mouse                           ------------------------------------------------------------   0
human                           ------------------------------------------------------------   0
mosquito                        S--EVSNKFSTVTAVAAASLNNNVV----------------VS---------QKPILT--   149
Dmelanogaster                   SGNSAASSYSETGAVSLAMVSGSTNLYSAYQRSQT-TDNTQSNQQHVVNSAENMPVIIKK   150
Sexual_Dmercatorum              NSLSAASSYSESSAVPPAFVSGSTLHIKRELEDEVQLEEVHDQDQDNDSNSENCPVNSNS   163
parthenogenetic_Dmercatorum     NSLSAASSYSESSAVPPAFVSGSTLHIKRELEDEVQLEEVHDQDQDNDSNSENCPVNSN-   161

mouse                           ------------------------------------------------------------   0
human                           ------------------------------------------------------------   0
mosquito                        --------------------------PANTSAMNINNNLTAKMATVKQQIPAGRSLLISSR   184
Dmelanogaster                   ELADLDYTVCQKRLRLSGGDKKSQ------------------IQDEVHLIPPGGSLLRKRN   193
Sexual_Dmercatorum              GSNSSGIRKRTNSCRSTGGSSKVAAATATPTISIPSRMIHRDPVIEPYIPPGGSLLRKSN   223
parthenogenetic_Dmercatorum     ---SSGIRKRTNSCRSTGGSSKVAAATATPTISIPSRMIHRDPVIEPYIPPGGSLLRKSN   218
                             4 aa deletion

mouse                           -------------------MDFLWALET--------PQTATTM-----------------   16
human                           -------------------MDFFRVVEN--------QQPPATM-----------------   16
mosquito                        IKQQMNRIPTVST---S---DFLR--ERETAVPLHRPDTPLSL--DEDPPEFKHNIDLAT   234
Dmelanogaster                   NQDIIRKSGELSG---SDSIKY------------QRPDTPHSLTDEVAASEFRHNVDLRA   238
Sexual_Dmercatorum              TQHKLQQQKLQQQQQQQQRLTYLLSSNNYNNSNNNSNNNSYSMPEDEVLPVFRHNVDLRA   283
parthenogenetic_Dmercatorum     TQHKLQQQKLQQQQQQQQRLTYLLSSNNYNNSNNNSNNNSYSMPEDEVLPVFRHNVDLRA   278
                                                     :               :   ::

mouse                           ----PLNVNFTNRNYDLDYDS-----------------------VQPY------------   37
human                           ----PLNVSFTNRNYDLDYDS-----------------------VQPY------------   37
mosquito                        CTIGSNRLSLTGHSRHYKNHQSHHDDPSSHRIINMLKEHLEDNESSSFRTCMASSTGEVG   294
Dmelanogaster                   CVMGSNNISLTGNDSDVNY-------------IKQISRELQNTGKDPLPVR-YIP-----   279
Sexual_Dmercatorum              CVMGSNNISLTN-SSDANI-------------IDLLSRELQNTSKERIDLP-YRIPGDPP   328
parthenogenetic_Dmercatorum     CVMGSNNISLTN-SSDANI-------------IDLLSRELQNTSKERIDLP-YRIPGDPP   323
                                      .:.:*.  .  .  .                        .

mouse                           FICDEEENFYHQQQQSE---------LQPPAPSEDIWKKFELLPTPPLSPSRRSGLCSPS   88
human                           FYCDEEENFYQQQQQSE---------LQPPAPSEDIWKKFELLPTPPLSPSRRSGLCSPS   88
mosquito                        SLTDLLLNDLEEMEEM--------------------------------ES-R----DGDD   316
Dmelanogaster                   PINDVLDVLNQHSNSTGGQQQLNQQQLDEQQQAIDIATGRNTVDSPPTTG-S----DSDS   334
Sexual_Dmercatorum              IITDVLEVLNQQAQQSASSAA----------AAAAAAAAAATLSPPATTA-T----SSDS   373
parthenogenetic_Dmercatorum     IITDVLEVLNQQAQQSASSAA----------AAAAAAAAAATLSPPATTA-T----SSDS   368
                                   *  : : .  :                                          . .

mouse                           YVAVATSFSPRE--------DDDGGGG--NFSTADQLEMM--TELLGGDMVNQSFICDPD   136
human                           YVAV-TPFSLRG--------DNDGGGG--SFSTADQLEMV--TELLGGDMVNQSFICDPD   135
mosquito                        SHGEELDSDTDSNADSSSRSSSKGGGISGGYTHAHNQEMSPSSSSSSSSYEYQGTHVG   376
Dmelanogaster                   DDGEPLNFDLRHH------RTSKSGSNASITTNNNNSN--NKNNKLKNNSNGMLHMMHIT   386
Sexual_Dmercatorum              D------------------------------------SD--YGDCSMGESSCSASIMRHIS   396
parthenogenetic_Dmercatorum     D------------------------------------SD--YGDCSMGESSCSASIMRHIS   391
                                                                      :           .        .

                                                  Myc box 3
mouse                           DETFIKNIIIQDCMWSGFSAAAKLVSEKL--ASYQAARKDSTSLSPARGHSVCST-----   189
human                           DETFIKNIIIQDCMWSGFSAAAKLVSEKL--ASYQAARKDSGSPNPARGHSVCST-----   188
mosquito                        DHSYTRPKARYNLAELGVQTPSDSEDEEIDVVSIGE-KNLPTNPTPRDKRHVESRVALKI   435
Dmelanogaster                   DHSYTRCNDMVD-DGPNLETPSD-SDEEIDVVSYTD-KKLPTNPSCHLMGALQFQMAHKI   443
Sexual_Dmercatorum              DHSYTRCNEVE----ANLDTPSD-SDEEIDVVSLND-KKLPTNPSDRDRRVLQTKVANKI   450
parthenogenetic_Dmercatorum     DHSYTRCNEVE----ANLDTPSD-SDEEIDVVSLND-KKLPTNPSDRDRRVLQTKVANKI   445
                                *.:: :            ...: :.  .*::  .*    ::   .  .       :
```

```
mouse                           -----------SSLYLQDLTAAASECIDP--SVVFPYPLNDSSSPKSCTSSDSTAFSPSS     236
human                           -----------SSLYLQDLSAAASECIDP--SVVFPYPLNDSSSPKSCASQDSSAFSPSS     235
mosquito                        RKHPQGNPSHHH--------------RRRHSGEDYPSHHGMSSSSSQHSPSKSYGYSPNY     481
Dmelanogaster                   SIDHMK-QKPRYNNFNLPYTPASSSPVKSVANSRYPSPSST---PYQNCSSASPSYSPLS     499
Sexual_Dmercatorum              SSDNRIVAHRSSRRYELPYTPASSSPVKSVANSRYPSPSST---PYQGAATGPATYSPES     507
parthenogenetic_Dmercatorum      SSDNRIVAHRSSRRYELPYTPASSSPVKSVANSRYPSPSST---PYQGAATGPATYSPES    502
                                                               .  :*   .      .        :**

mouse                           DSLLSS-ESS----------------------------------------------PRAS     249
human                           DSLLSSTESS----------------------------------------------PQGS     249
mosquito                        LTPASSTSIS----------------G---------------SNTPLPPNSSSISNPR--     508
Dmelanogaster                   VDSSNVSSSSSSSSSSQSSFTTSSSNKGRKRSSLKDPGLLISSSSVYLPGVNNKVTH----     555
Sexual_Dmercatorum              SSSSSDCTTP-------SIALGVGAGGKK-----------NRKPFYMPDCNDDLLTAKRQ     549
parthenogenetic_Dmercatorum     SSSSSDCTTP-------SIALGVGAAGKK-----------NRKPFYMPDCNDDLLTAKRQ     544
                                    .                    G526A

mouse                           PEPLVLHEETPPTTSSDSEEEQEDEEEIDVVSVEKRQTPAKRSESGSSPS--RGHSKPPH     307
human                           PEPLVLHEETPPTTSSDSEEEQEDEEEIDVVSVEKRQAPGKRSESGSPSA--GGHSKPPH     307
mosquito                        ---------KR-----------------------------------------------PSK     513
Dmelanogaster                   ------------------------------SSMMSKKSRGK-KVVGTSSGNTSPIS-SGQ     583
Sexual_Dmercatorum              PRGYLLSKKRPLKRTHYSSYGF-DAKE--VRSVLSHASNSV-STIGSSSS---NSSKSGH     602
parthenogenetic_Dmercatorum     PRGYLLSKKRPLKRTHYSSYGF-DAKE--VRSVLSHASNSV-STIGSSSSNSSNSSKSGH     600
                                                                         3 aa addition     :

mouse                           SPLVLKRCHVSTHQHNYAAPPSTRKDYPAAKRAKLDSGRVLKQISNNR----KCSSPRSS     363
human                           SPLVLKRCHVSTHQHNYAAPPSTRKDYPAAKRVKLDSVRVLRQISNNR----KCTSPRSS     363
mosquito                        DDRSKNRH----HQH------------------RNKKQRIPG-------KTIARSPESSE     544
Dmelanogaster                   DVDAMDRN----WQR------------------R--SGGIATSTSSNSSVHRKDFVLGFD     619
Sexual_Dmercatorum              SNG----S----HSS------------------N--SGHSNGSISNGSGINSLKRHLSID     634
parthenogenetic_Dmercatorum     SNG----S----HSS------------------N--SGHSNGSISNGSGINSLKRHLSID     632
                                .            .                   .  .                      .
                                           Helix-loop-helix DNA-binding domain
mouse                           DTEENDKRRTHNVLERQRRNELKRSFFALRDQIPELENNEKAPKVVILKKATAYILSIQA     423
human                           DTEENVKRRTHNVLERQRRNELKRSFFALRDQIPELENNEKAPKVVILKKATAYILSVQA     423
mosquito                        EQETLEKRNLHNDMERQRRIGLKNLFEELKRQIPNLRDKERAPKVNILREAAVLCTRLNR     604
Dmelanogaster                   EADTIEKRNQHNDMERQRRIGLKNLFEALKKQIPTIRDKERAPKVNILREAAKLCIQLTQ     679
Sexual_Dmercatorum              EADTIEKRNLHNDMERQRRIGLKNLFEALKTQIPNIRDKERAPKVNILREAARLCEQLTS     694
parthenogenetic_Dmercatorum     EADTIEKRNLHNDMERQRRIGLKNLFEALKTQIPNIRDKERAPKVNILREAARLCEQLTS     692
                                : :   **. ** :*****  **. *  *: *** :.::*:**** **::*:     :

mouse                           DEHKLTSEKDLLRKRREQLKHKLEQLRNSGA--------------     454
human                           EEQKLISEEDLLRKRREQLKHKLEQLRNSCA--------------     454
mosquito                        EQEQLNALRKQ----QQRLYARVRQLRTSLHTQ-------RRVMD     638
Dmelanogaster                   EEKELSMQRQL-------LSLQLKQRQDTLASYQMELNESRSVSG     717
Sexual_Dmercatorum              EERDLNVKRQL-------LKAKLKQQQEQLARMRLNLSKNE----     728
parthenogenetic_Dmercatorum     EERDLNVKRQL-------LKAKLKQQQEQLARMRLNLSKNE----     726
                                ::..*   ..        *  ::.* :
```

**Figure S8:**

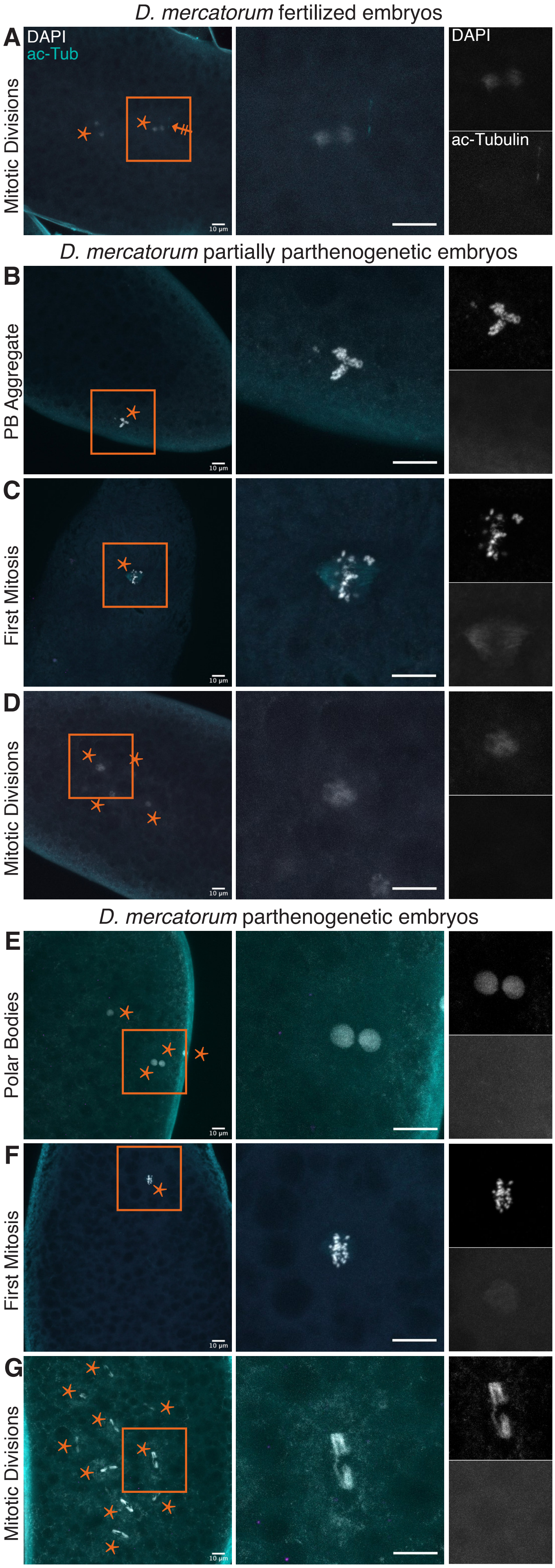

D. mercatorum fertilized embryos
A
DAPI
ac-Tub
Mitotic Divisions
DAPI
ac-Tubulin
10 µm
D. mercatorum partially parthenogenetic embryos
B
PB Aggregate
C
First Mitosis
D
Mitotic Divisions
D. mercatorum parthenogenetic embryos
E
Polar Bodies
F
First Mitosis
G
Mitotic Divisions


**Figure S9:**

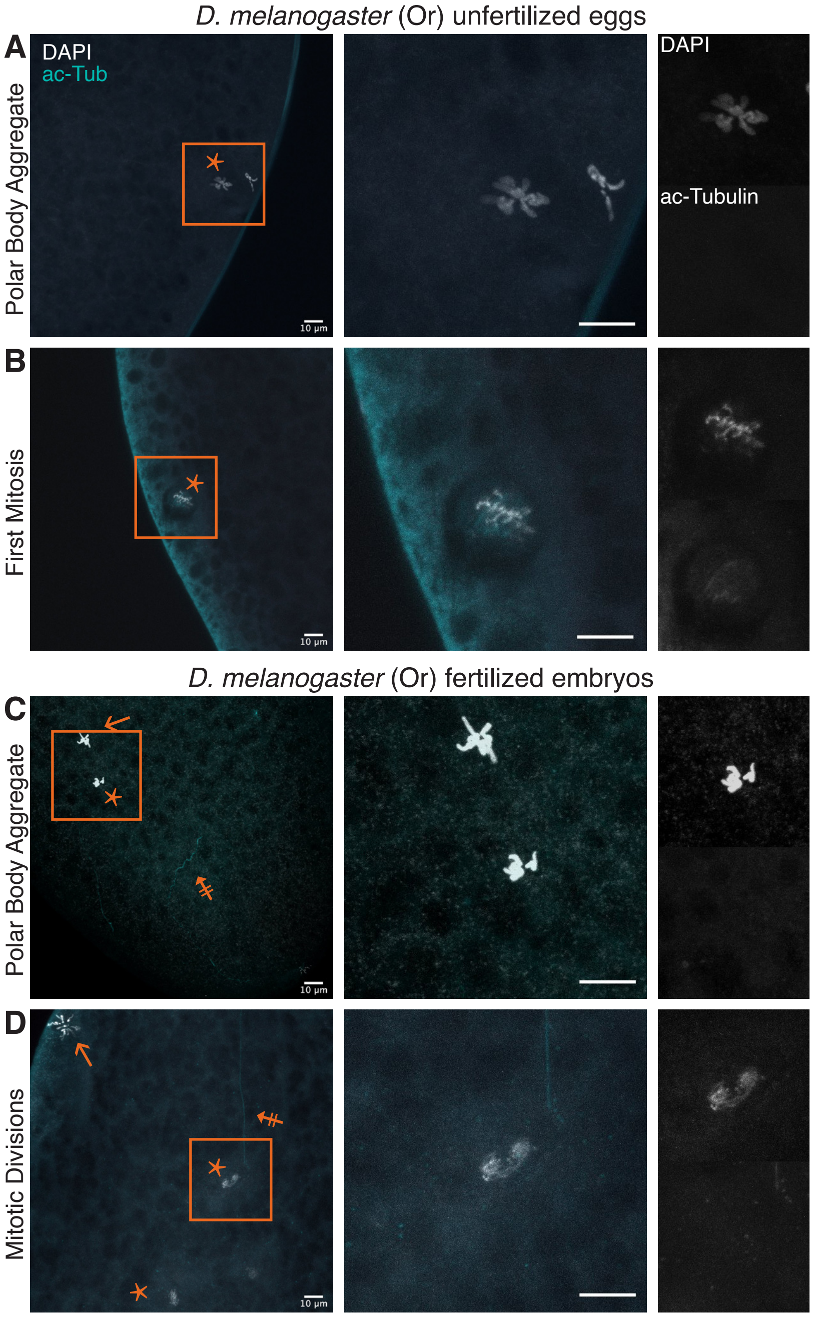

D. melanogaster (Or) unfertilized eggs
A
DAPI
ac-Tub
Polar Body Aggregate
DAPI
ac-Tubulin
B
First Mitosis
D. melanogaster (Or) fertilized embryos
C
Polar Body Aggregate
D
Mitotic Divisions
10 μm


**Figure S10:**

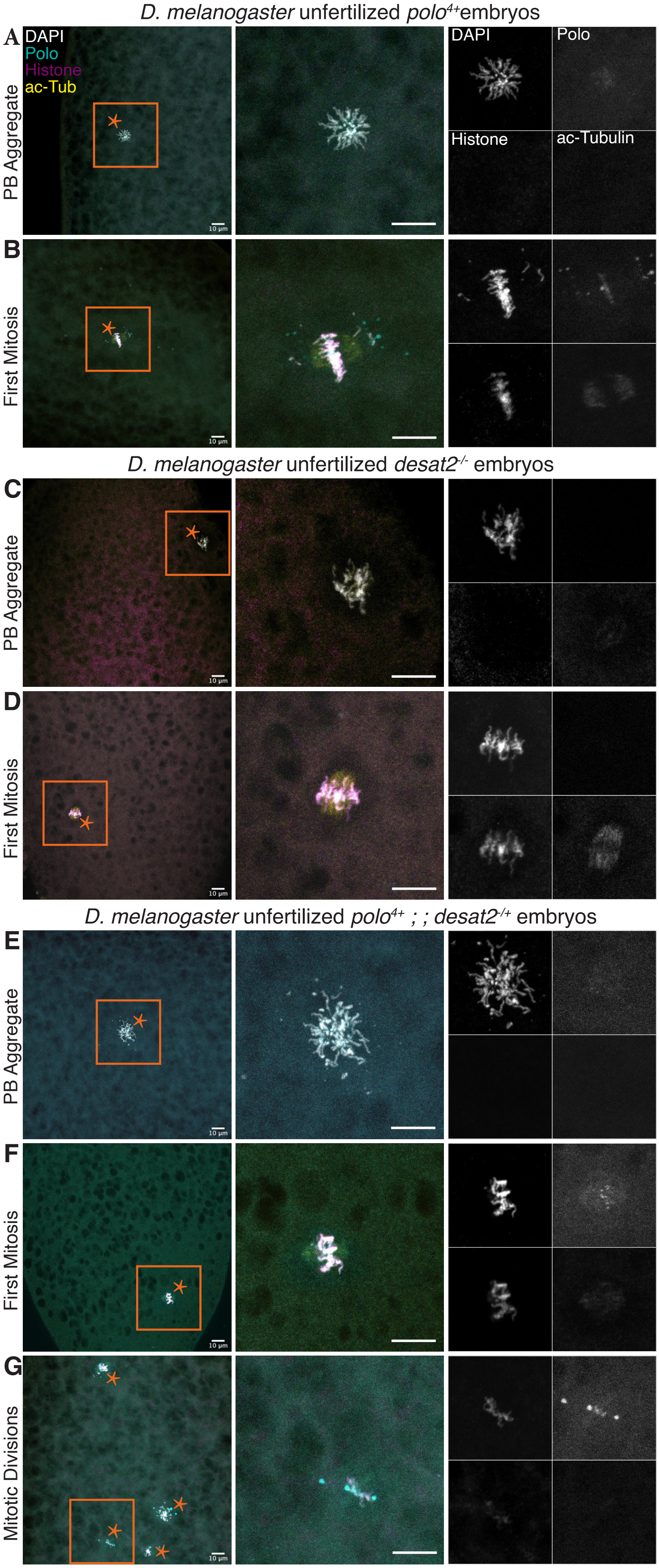

D. melanogaster unfertilized polo4+embryos
A
DAPI
Polo
Histone
ac-Tub
PB Aggregate
DAPI
Polo
Histone
ac-Tubulin
B
First Mitosis
D. melanogaster unfertilized desat2-/- embryos
C
PB Aggregate
D
First Mitosis
D. melanogaster unfertilized polo4+ ; ; desat2-/+ embryos
E
PB Aggregate
F
First Mitosis
G
Mitotic Divisions


**Figure S11:**

| Gene | Function | Screened with | Percent | *p* value |
|---|---|---|---|---|
| *ana2* | Centriole | CRISPR | 0 | |
| *asl* | Centriole/PCM | CRISPR/ubiquitous expression | 0-0.1% | 0-0.50 |
| *cnn* | PCM | mutant | 0.1% | 0.38 |
| *cyclinE* | Cell cycle | UAS/Gal4 overexpression | 0 | |
| *morula* | Cell cycle | CRISPR | 0-0.1% | 0-0.40 |
| *Plk4* | Centriole/ Centrosome | CRISPR/UAS/Gal4 overexpression | 0 | |
| *Plp* | PCM | CRISPR | 0.1% | 0.24 |
| *plu* | Translation | CRISPR | 0.1% | 0-0.50 |
| *png* | Translation | CRISPR | 0 | |
| *polo* | Cell cycle | mutants/endogenous promotor overexpression | 0-0.1% | 0-0.49 |
| *Sas-6* | PCM | CRISPR/ubiquitous expression | 0-0.1% | 0-0.40 |
| *slimb* | SCF complex/ Cell cycle | CRISPR/mutant | 0-0.3% | 0-0.17 |
| *Rca1* | Cell cycle | UAS/Gal4 overexpression | 0 | |
| *tefu /atm* | Serine/threonine kinase | RNAi/mutant | 0 | |

**Table S1**

| Gene | Function | Screened with | Percent |
| --- | --- | --- | --- |
| *CG3436* | Cell cycle | mutant | 0 |
| *dhd* | embryonic development | mutant | 0 |
| *Klp64D* | Motor protein | RNAi/mutant | * |
| *Trx-2* | Redox | RNAi/mutant | * |
| *w* | Eye pigment transporter | RNAi/mutant | * |

**Table S2**

| *Drosophila* species/genotype | Successful F2 | No F2 |
|---|---|---|
| ***melanogaster*** | | |
| *polo*$^{4+}$ *; desat1*$^{-/+}$ | 1 | 0 |
| *polo*$^{4+}$ *; desat2*$^{-/+}$ | 2 | 0 |
| ***mercatorum*** | | |
| partially parthenogenetic | 17 | 7 |

**Table S3:**
